# *Plasmodium falciparum* Calcium Dependent Protein Kinase 4 is critical for male gametogenesis and transmission to the mosquito vector

**DOI:** 10.1101/2021.06.06.447280

**Authors:** Sudhir Kumar, Meseret T. Haile, Michael R. Hoopmann, Linh T. Tran, Samantha A. Michaels, Seamus R. Morrone, Kayode K. Ojo, Laura M. Reynolds, Ulrike Kusebauch, Ashley M. Vaughan, Robert L. Moritz, Stefan H.I. Kappe, Kristian E. Swearingen

## Abstract

Gametocytes of the malaria parasite *Plasmodium* are taken up by the mosquito vector with an infectious blood meal, representing a critical stage for parasite transmission. Calcium dependent protein kinases (CDPKs) play key roles in calcium-mediated signaling across the complex life cycle of the parasite. We sought to understand their role in human parasite transmission from the host to the mosquito vector and thus investigated the role of the human-infective parasite *Plasmodium falciparum* CDPK4 in the parasite life cycle. *P. falciparum cdpk4*^−^ parasites created by targeted gene deletion showed no effect in blood stage development or gametocyte development. However, *cdpk4*^−^ parasites showed a severe defect in male gametogenesis and the emergence of flagellated male gametes. To understand the molecular underpinnings of this defect, we performed mass spectrometry-based phosphoproteomic analyses of wild type and *Plasmodium falciparum cdpk4*^−^ late gametocyte stages, to identify key CDPK4-mediated phosphorylation events that may be important for the regulation of male gametogenesis. We further employed *in vitro* assays to identify these putative substrates of *Plasmodium falciparum* CDPK4. This indicated that CDPK4 regulates male gametogenesis by directly or indirectly controlling key essential events such as DNA replication, mRNA translation and cell motility. Taken together, our work demonstrates that PfCDPK4 is a central kinase that regulates exflagellation, and thereby is critical for parasite transmission to the mosquito vector.

**IMPORTANCE:** Transmission of the malaria parasite to the mosquito vector is critical for the completion of the sexual stage of the parasite life cycle and is dependent on the release of male gametes from the gametocyte body inside the mosquito midgut. In the present study, we demonstrate that PfCDPK4 is critical for male gametogenesis and is involved in phosphorylation of proteins essential for male gamete emergence. Targeting PfCDPK4 and its substrates may provide insights into achieving effective malaria transmission-blocking strategies.

## INTRODUCTION

The malaria parasite *Plasmodium falciparum* (Pf) remains a major causative agent of mortality and morbidity in developing countries across the world. Pf is an obligate intracellular parasite whose life cycle alternates between a human host and an arthropod *Anopheles* mosquito vector. In the human host, the parasite replicates asexually within red blood cells and develops over a ~48 hr cycle as ring, trophozoite and schizont stages. Some asexually replicating parasites can enter a terminally differentiated sexual developmental pathway known as gametocytogenesis, which culminates with the formation of mature, transmissible gametocytes. Factors responsible for gametocyte induction include stress factors such as exposure to drugs and availability of host serum factors such as phosphatidylcholine and phosphatidylethanolamine (1, 2). Gametocytes become activated upon uptake by the mosquito vector and form gametes (female; macrogametes and male; microgametes). While the male gametocyte undergoes significant changes in morphology and size reduction resulting from three rapid rounds of mitosis to form eight flagellar microgametes, the female gametocyte forms a single macrogamete. Fusion of male and female gametes leads to the formation of a zygote, which in turn develops into motile ookinete. Ookinetes penetrate the mosquito midgut epithelium to develop into oocyst stages which eventually produce transmissible sporozoites (3).

Gametogenesis within the mosquito midgut may be triggered by a combination of factors including drop in temperature (4, 5), increase in pH (5), and/or exposure to xanthurenic acid (XA), a metabolite of tryptophan (6, 7). Gametogenesis is also linked to mobilization of intracellular calcium (Ca^2+^) stores, which can regulate Ca^2+^ dependent protein function (8, 9). The Pf genome encodes seven distinct CDPKs, which are key Ca^2+^ effectors in the parasite that have roles throughout the *Plasmodium* life cycle (10). The fact that CDPKs are not encoded by the human genome makes them attractive drug targets. CDPK4 in particular has been implicated in playing an essential role in gametocyte transmission. Studies in the rodent malaria model *Plasmodium berghei* (Pb) identified PbCDPK4 (PBANKA_0615200) as the Ca^2+^ effector in gametocytes, and activated male gametocytes lacking PbCDPK4 failed to undergo exflagellation (8). The role of PfCDPK4 (PF3D7_0717500) in transmission of Pf parasites to the mosquito vector however has only been studied via pharmacological inhibition with bumped kinase inhibitors (11, 12) which may affect other kinases in the parasite.

Here, we used Clustered Regularly Interspaced Short Palindromic Repeats-Cas9 (CRISPR-Cas9)-based gene deletion to assess PfCDPK4 function. We demonstrate that PfCDPK4 is dispensable for asexual replication, sexual stage commitment and gametocytogenesis, but is essential for male gametogenesis, the emergence of flagellated male gametes and thereby transmission to the mosquito vector. Further, we used quantitative phosphoproteomics to gain insights into the phosphosignaling network of PfCDPK4, including identification of putative substrates.

## RESULTS

### PfCDPK4 is expressed in the asexual and sexual stages of the parasite

To analyze expression of PfCDPK4, antisera was generated against a synthetic KLH-conjugated peptide (KMMTSKDNLNIDIPS) designed from the J-domain (Fig. 1A). Indirect immunofluorescence assays (IFAs) done on thin blood smears of in vitro cultured PfNF54 revealed that PfCDPK4 is abundantly expressed in the ring, trophozoite and schizont stages as well as in free merozoite stages (Fig. 1B). PfCDPK4 expression was also detected in gametocytes from stage II through stage V (Fig. 2A). Counterlabelling with male (anti-tubulin II) or female (anti-Pfg377) gametocyte specific antibodies revealed that PfCDPK4 is expressed in both male and female gametocytes (Fig. 2A and 2B). Previous studies have shown that stage-specific *PbCDPK4* conditional knockout (cKO) parasites with highly reduced levels of PbCDPK4 expression in sporozoite stages display a decrease in infectivity for hepatocytes and thus establish a role for PbCDPK4 in hepatocyte invasion (13). We thus further analyzed PfCDPK4 expression in sporozoites and observed a circumferential localization in a similar fashion to circumsporozoite protein (PfCSP) (Fig. 2C).

**Figure 1.**
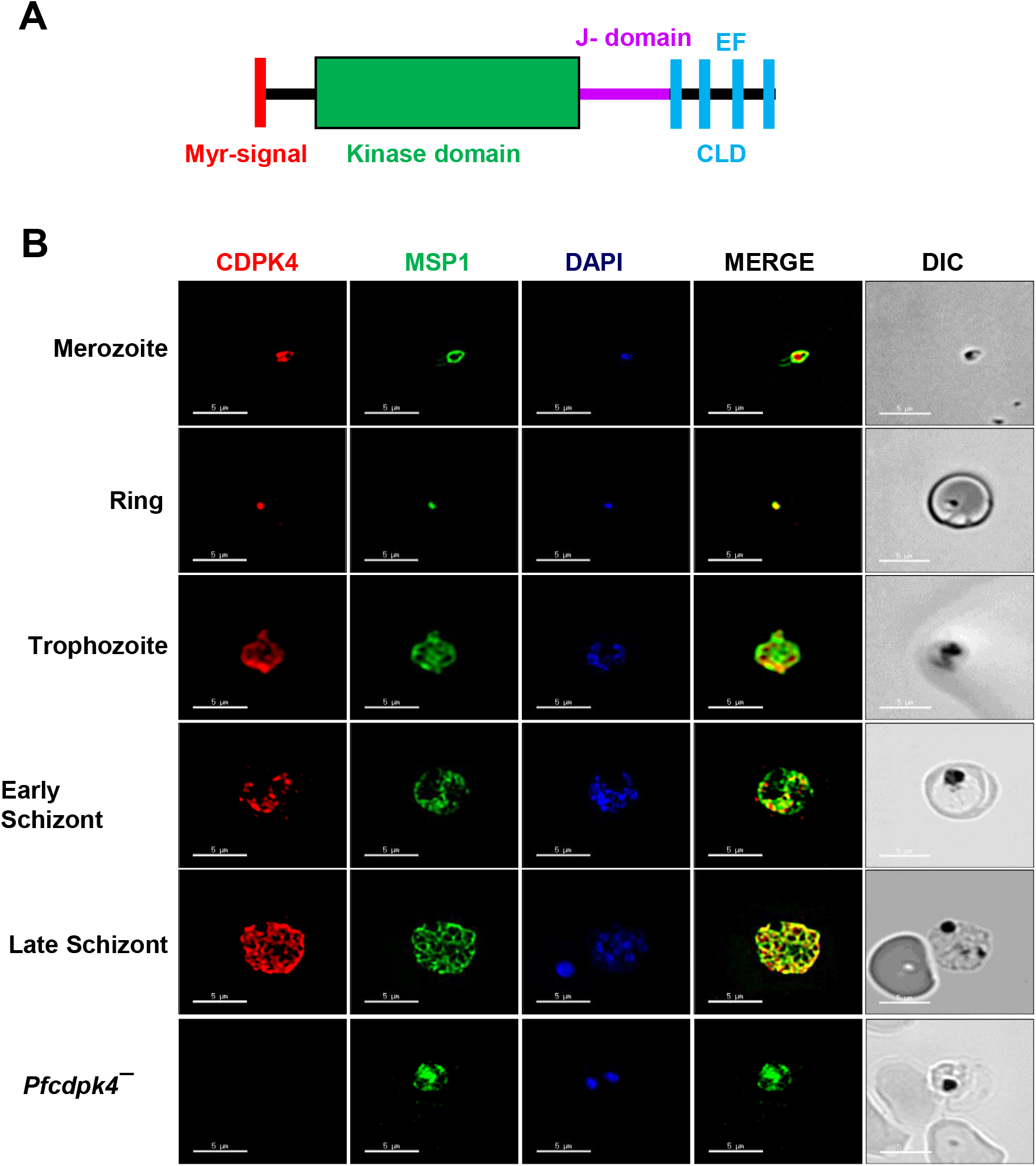
Expression and localization of PfCDPK4 in asexual sages of parasite. (**A**) Schematic for PfCDPK4 showing an N-terminal myristoylation signal (red) followed by a kinase domain (green), a junctional domain (J-domain) (pink), and a C-terminal calmodulin-like domain (CLD) containing four EF-hand motifs (cyan). (**B**) Immunofluorescence assays were performed on WT NF54 asexual blood stages (merozoite, ring, trophozoite, early and late schizonts) and *Pfcdpk4^−^* trophozoites to colocalize PfCDPK4 (red) in combination with merozoite surface protein 1 (MSP1, green). The parasite nucleus was localized with 4′,6-diamidino-2-phenylindole (DAPI) (in blue). Scale bar = 5 μm.

**Figure 2.**
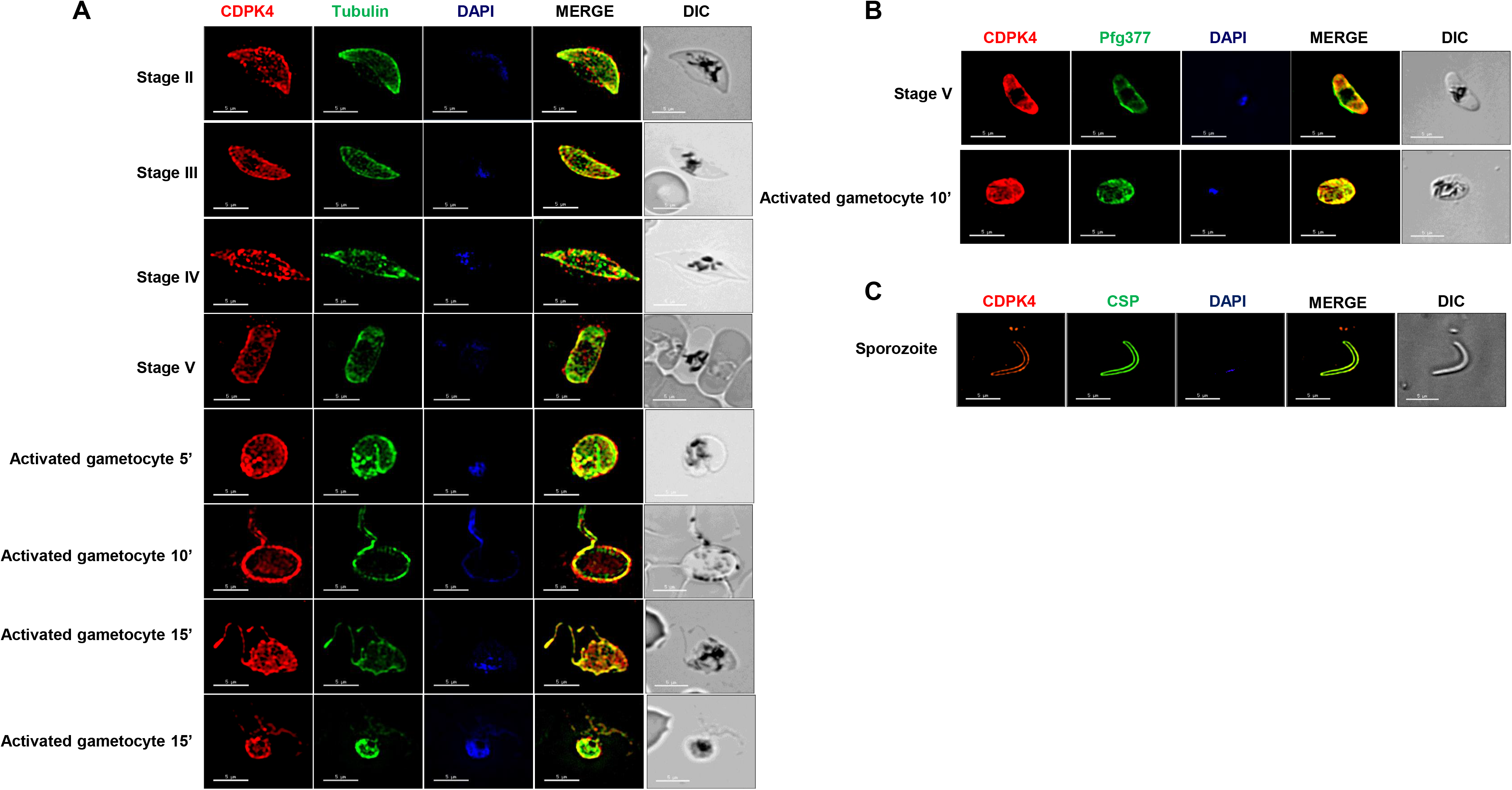
Expression and localization of PfCDPK4 in sexual stages of parasite. (**A**) Immunofluorescence assays were performed on WT NF54 sexual (stage II-V gametocytes) and 10 min post activation using thin smears and anti-PfCDPK4 antisera (in red) either in combination with α-Tubulin II (male gametocytes) (**B**) Immunofluorescence assays were performed on stage V female gametocytes and 10 min post activation using thin smears and anti-PfCDPK4 antisera (in red) in combination with anti-Pfg377 (marker for female gametocytes, in green). (**C**) Immunofluorescence assays were performed on sporozoites for the expression of PfCDPK4 (in red) and PfCSP (in green). Parasite nucleus was stained with DAPI (blue). Scale bar = 5 μm.

### Disruption of the *PfCDPK4* has no effect on intra-erythrocytic parasite development

For functional analysis, the endogenous *PfCDPK4* gene was disrupted using CRISPR/Cas9 (Fig. 3A). Gene deletion parasites (*Pf cdpk4^−^*) were confirmed by a set of diagnostic PCRs with oligonucleotides specific for the *PfCDPK4* locus and its upstream (5’) and downstream (3’) regions (Fig. 3A, 3B and 3C). Two clones for *Pf cdpk4^−^* parasites (clone IC2 and ID2) were used for phenotypic analysis. To analyze the role of PfCDPK4 in asexual parasite stages, a growth rate experiment was set up using *Pf cdpk4^−^* parasites (clone IC2 and ID2) along with wildtype (WT) NF54 parasites. Growth was monitored over two replication cycles. Giemsa-stained thin smears prepared every 48-hrs from the culture indicated that the growth rate of *Pf cdpk4^−^* parasites was similar to WT parasites (Fig. 4A).

**Figure 3.**
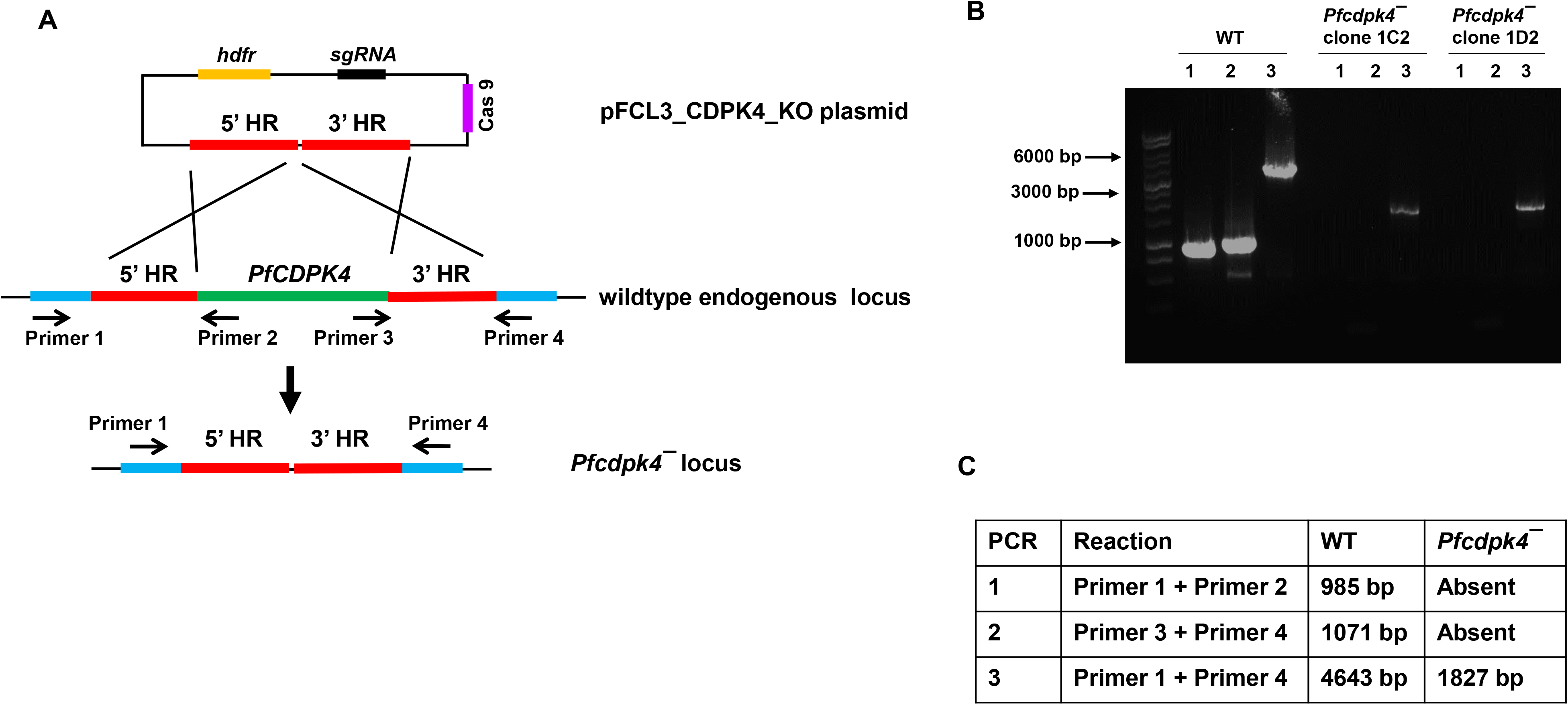
Disruption of the PfCDPK4 locus via CRISPR/Cas9. The schematic shows the strategy for deleting *PfCDPK4*. The pFCL3_CDPK4_KO plasmid has homology regions 5’ (5’HR) and 3’ (3’HR) not of the *PfCDPK4* locus, a guide RNA sequence (sgRNA) and human dihydrofolate reductase (hDHFR) locus and Cas 9 cloned. (**B**) Confirmation of *PfCDPK4* deletion by diagnostic PCR. The oligonucleotides were designed from outside 5’HR and 3’HR and *PfCDPK4* locus and positions are indicated by arrows in (**A**). The expected sizes for different set of PCRs are indicated in (**C**).

**Figure 4.**
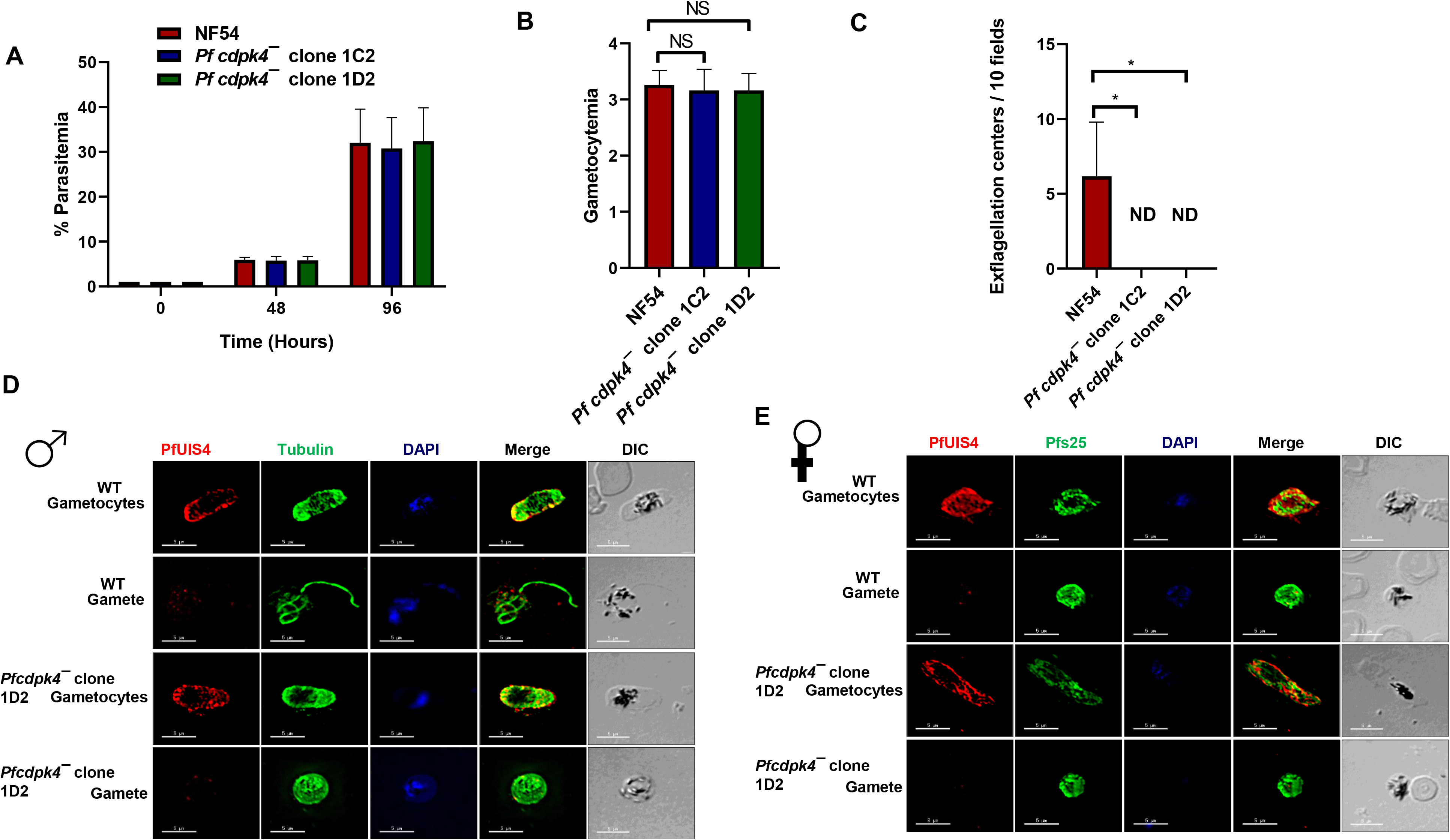
*Pfcdpk4^−^* blood stages show no phenotype and undergo gametocytogenesis. (**A**) Ring stage synchronous cultures for WT and two clones of *Pfcdpk4^−^* (clone 1C2 and 1D2) were plated to measure parasite growth over the course of two erythrocytic cycles. Total parasitemia was determined by counting the parasites from Giemsa-stained thin blood smears. Data were averaged from three biological replicates and presented as the mean ± standard deviation (SD). ns, not signifcant (unpaired two-tailed Student’s t test. (**B**) Ring stage synchronous cultures for WT and two different clones of *Pfcdpk4^−^* (clone 1C2 and 1D2) were tested for their potential to form gametocytes. Gametocytemia was measured on day 15 using Giemsa stained smears. Data were averaged from three biological replicates and presented as the mean ± standard deviation (SD). (**C**) Number of exflagellation centers (mitotic division during male gametogenesis) per field at 15 min post-activation. Data were averaged from three biological replicates and presented as the mean ± standard deviation (SD). (**D**) and (**E**) IFAs performed on thin blood smears of mature stage V gametocytes activated for 20 mins *in vitro* for WT or *Pfcdpk4^−^* (clone 1D2) and were stained for α-tubulin II (green), a male-specific marker, and Pfg377 (red), a marker for female gametes in an IFA. Anti-PfUIS4 was used to stain parasitophorous vacuolar membrane. α-Tubulin II staining showed male gametes emerging from an exflagellating male gametocyte in the WT parasite. The *Pfcdpk4^−^* gametocytes were defective for male gametocyte exflagellation. Female gametes did not show any defect in egress from gametocyte body.

### *Pf cdpk4^−^* parasites undergo gametocytogenesis but fail to form male gametes

We next analyzed the ability of *Pf cdpk4^−^* parasites to generate gametocytes. Gametocytemia was scored for all the cultures on day 15 of *in vitro* culture using Giemsa stained culture smears and microscopic inspection. *Pf cdpk4^−^* parasites were able to develop into mature stage V gametocytes and had similar gametocytemia compared to WT parasites (Fig. 4B). We further tested whether *Pf cdpk4^−^* gametocytes undergo gametogenesis. Day 15 gametocyte cultures for WT and *Pf cdpk4^−^* were activated by addition of O^+^ human serum and dropping the temperature from 37°C to room temperature (RT). Activated gametocytes were used to prepare a temporary live wet mount of cultures and exflagellation centers were measured in 10 random fields of microscopic view at 40x magnification. Intriguingly, we did not observe any exflagellation centers for *Pf cdpk4^−^* (Fig. 4C) indicating an exflagellation defect. To confirm this defect, IFAs were performed by making thin culture smears for WT and *Pf cdpk4^−^* activated gametocytes 10 min post activation and parasites were stained with anti-tubulin antibody. Lack of observable release of male gamete exflagella from the gametocyte body confirmed an exflagellation defect in *Pf cdpk4^−^* (Fig. 4D). Female *Pf cdpk4^−^* gametes were stained with Pfs25 antibody and appeared similar to WT gametes (Fig. 4E). These results indicate *Pf* CDPK4 is critical only for male gametogenesis.

We next examined the transmissibility of *Pf cdpk4^−^* gametocytes to female *Anopheles stephensi* mosquitoes. Infectious blood meals of WT and *Pf cdpk4^−^* stage V gametocytes were prepared using standard methods and fed to mosquitoes. Mosquito midguts were dissected on Day 7 post feed which revealed that *Pf cdpk4^−^* parasites displayed a complete absence of oocysts (Fig. 5) in comparison to well-infected WT controls. Taken together, these results reveal that PfCDPK4 is indispensable for transmission to the mosquito vector via a critical function in male gametogenesis.

**Figure 5.**
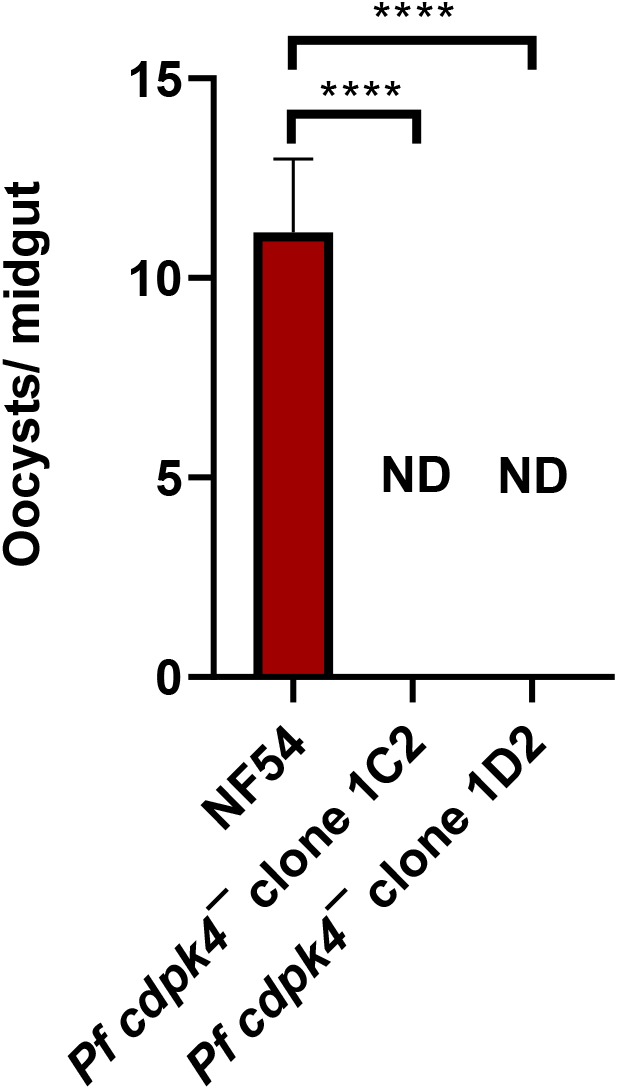
The *Pfcdpk4^−^* parasites do not establish infection in mosquitoes. Mosquitoes were dissected on day 7 post feed and number of oocysts were measured per midgut. Data were averaged from three biological replicates with a minimum of 50 mosquito guts and presented as the mean ± standard deviation (SD).

### Comparative phosphoproteomics elucidates the PfCDPK4 signaling network

In order to dissect the phosphorylation regulatory networks via which PfCDPK4 might be functioning and to determine which cellular processes it might be regulating in the parasite, we performed a comparative phosphoproteomic analysis of WT and *Pf cdpk4^−^* parasites. Since *Pf cdpk4^−^* displays a defect in male gamete exflagellation (Fig. 4D), we chose to perform this analysis on mature stage V activated gametocytes. Gametogenesis was initiated by exposing parasites to xanthurenic acid (XA), a compound endogenous to the mosquito gut that induces release of intracellular Ca^2+^ stores (6, 7). We then employed a label-free quantitative mass spectrometry-based approach to detect protein residues that were hypophosphorylated in the *Pf cdpk4^−^* parasites when compared to WT parasites, thereby identifying phosphorylation events downstream of PfCDPK4 activation, including putative substrates of PfCDPK4. After application of strict criteria for confidence of peptide identification and phosphosite localization, this analysis identified a total of 8221 unique phosphosites on 9810 peptides from 1904 Pf proteins across both sample types (Supplementary Data S1). We next applied strict criteria to identify phosphosites that were significantly hypophosphorylated or absent in the *Pf cdpk4^−^* parasite, thereby representing putative substrates of PfCDPK4 or other kinases downstream of the phosphosignaling cascade initiated by the activity of PfCDPK4. In all, 1011 unique phosphosites on 1059 peptides from 542 Pf proteins met our criteria, including 193 phosphosites on 196 peptides from 170 proteins that were confidently quantified in WT parasites but not detected in *Pf cdpk4^−^* parasites (Supplementary Data S1).

In order to ascertain the putative function of proteins involved in the PfCDPK4-mediated phosphosignaling network, we performed gene ontology (GO) analysis on all proteins showing hypophosphorylation in *Pfcdpk4^−^* parasites. Enriched molecular function terms included histone binding, RNA-binding, mRNA-binding, and translation factor activity, i.e., regulation of protein transcription and translation (Fig. 6A). Other enriched terms include kinase and ATPase activity, indicating that PfCDPK4 is an upstream regulator of a phosphosignaling cascade. Indeed, phosphosites that were hypophosphorylated in *Pfcdpk4^−^* parasites were detected on 23 proteins with annotated or predicted protein kinase or protein phosphatase activity (Supplementary Data S2). Motif analysis of these hypophosphorylated phosphosites found that the most enriched motif was [K/R]XX[S/T] (Fig. 6B), i.e. the “Simple 1” motif, which is common to calcium-dependent kinases (14, 15).

**Figure 6.**
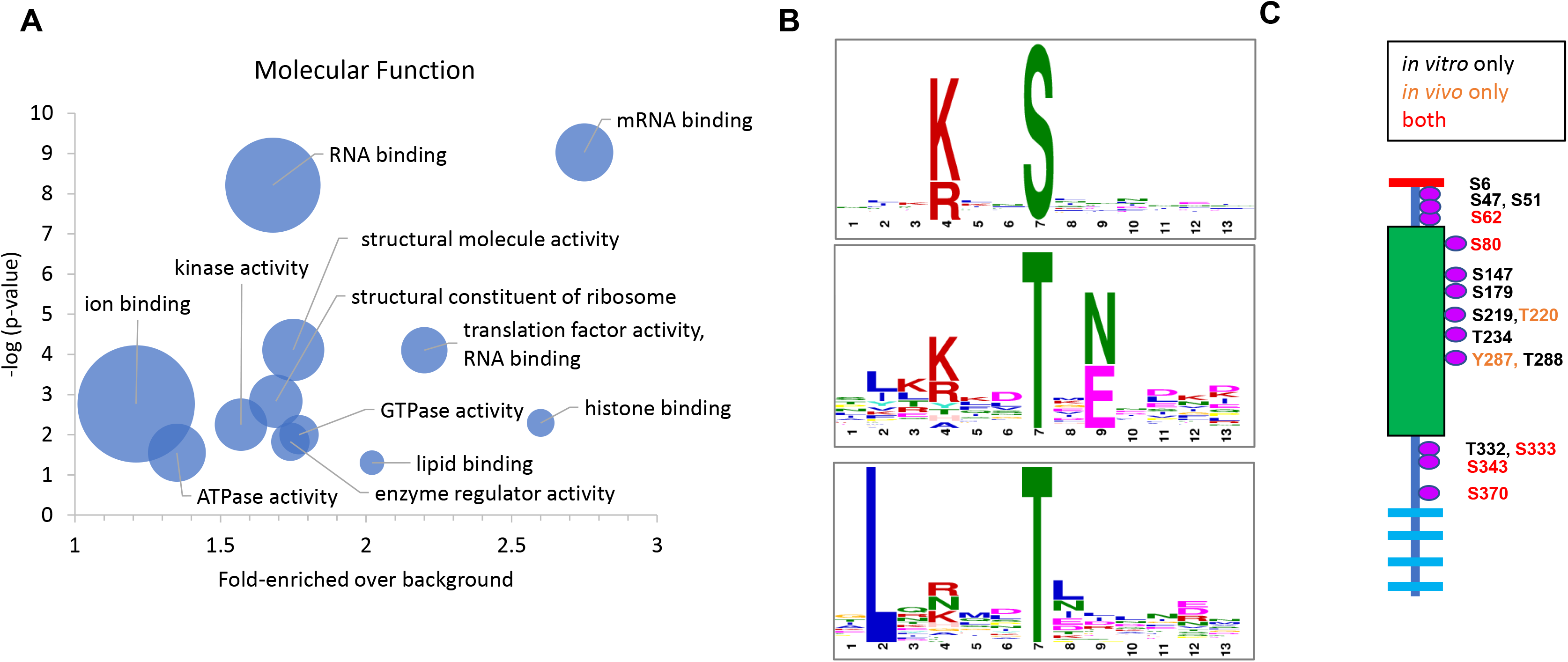
Phosphosignalling by PfCDPK4. (**A**) Molecular function terms enriched by gene ontology (GO) analysis of proteins bearing phosphosites that were hypophosphorylated in *Pfcdpk4^−^* gametocytes. The x-axis shows the magnitude of the enrichment of the term in the subset relative to its frequency in the genome. The y-axis is the negative log of the p-value showing the probability that the term is significantly enriched. The size of the bubble is proportional to the number of proteins in the subset described by the indicated GO term. (**B**) Motif analysis of phosphosites hypophosphorylated in *Pfcdpk4^−^* gametocytes revealed an enrichment for the Simple 1 motif [K/R]XX[S/T] favored by calcium dependent kinases, as well as a novel motif involving Leu at the −5 position. The phosphosite in each motif is at position 7 as indicated on the x-axis. The relative height of the letters is proportional to the bit score quantifying the likelihood of finding the represented residue at that position. (**C**) PfCDPK4 is extensively autophosphorylated *in vitro* (black and red text). Several of these sites were also observed *in vivo* in activated WT gametocytes (red text). The two phosphosites indicated with orange text were only observed *in vivo*.

### PfCDPK4 is phosphorylated *in vitro* and in the parasite

Autophosphorylation of PfCDPKs has been proposed to regulate their activity *in vitro* (16, 17). Phosphorylation of PfCDPK4 has been observed in asexual stages at S^80^ (18), but to date no phosphoproteomic studies of Pf gametocytes have been published, nor has autophosphorylation of PfCDPK4 been well-cataloged. *In vitro* studies with recombinant PfCDPK4 have shown that the kinase is activated when the presence of Ca^2+^ leads to autophosphorylation of the active site residue T^234^, and that absence of Ca^2+^ or a T234A mutation inactivates the kinase (17). In order to identify additional autophosphorylation sites for PfCDPK4, we performed an *in vitro* kinase assay in which recombinant PfCDPK4 was activated by the presence of CaCl_2_ in the reaction buffer, after which the protein was digested with trypsin and analyzed by LC-MS/MS. We identified a total of 14 phosphosites, including T^234^ (Fig. 6C). We also observed seven PfCDPK4 phosphosites in our phosphoproteomic analysis of WT parasites, of which five were among the autophosphorylation sites seen *in vitro*. The two phosphosites only observed *in vivo*, T^220^ and Y^287^ in the kinase domain, are adjacent to residues that were autophosphorylated in vitro, i.e. S^219^ and T^288^, which, in turn, were not observed *in vivo*. It is therefore possible that T^220^ and Y^487^ are PfCDPK4 autophosphorylation sites that were mislocalized in the interpretation of the identifying mass spectra, as opposed to representing the activity of other kinases modifying PfCDPK4.

### *In vitro* assays support identification of putative PfCDPK4 substrates

In order to further investigate putative PfCDPK4 substrates, we performed an LC-MS-based *in vitro* assay using recombinant PfCDPK4 and synthetic peptides bearing residues that we predicted to be phosphorylated by PfCDPK4 based on our phosphoproteomic analysis. The Pf proteins that we selected for testing include novel targets that we identified as well as Pf orthologues of proteins that have previously been annotated as putative substrates of CDPK4 (SOCs) based on work in *P. bergehi* (19). Synthetic peptides showing significant phosphorylation by recombinant PfCDPK4 *in vitro* support the identification five putative substrates of PfCDPK4: the uncharacterized protein PF3D7_0417600, PfCDPK1 (PF3D7_0217500), PfSOC3 (PF3D7_1440600), PfSOC7 (PF3D7_1437200; ribonucleoside-diphosphate reductase), and ATP-dependent 6-phosphofructokinase (PFK9; PF3D7_0915400) (Table 1, Supplementary Data S3, Supplementary Data S4).

**Table 1.**
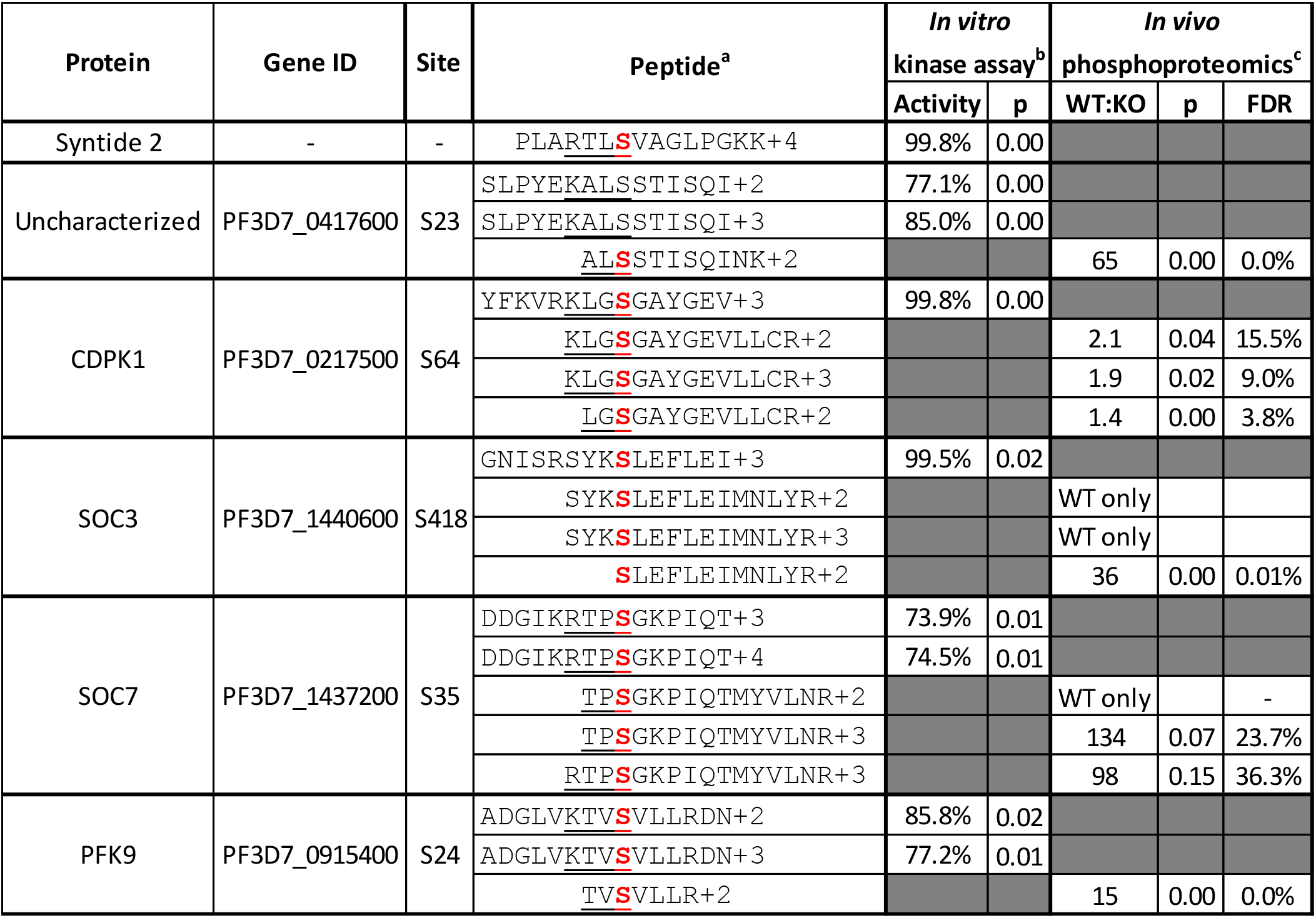
Putative PfCDPK4 substrates identified by an *in vitro* kinase assay. Synthetic peptides were reacted with recombinant PfCDPK4 and the extent of phosphorylation was quantified by LC-MS. Peptides confirming putative substrates of PfCDPK4 are shown. Extended data, including data from additional peptides assayed, are provided in Supplementary Data S3. (a) For each protein, peptide sequences are shown for the synthetic peptide assayed *in vitro* and for tryptic peptide observed *in vivo* from phosphoproteomic analyses of activated gametocytes. The charge state of the quantified peptide ion is indicated. Syntide 2 is a positive control standard peptide. The phosphorylated residue is shown in bolded red. The Simple 1 motif [K/R]XX[S/T] is underlined where present. The LC-MS/MS spectra for the phosphorylated version of the peptide SLPYEKALSSTISQI were of insufficient quality to confidently sequence the phosphopeptide or confirm which residue was phosphorylated. However, a peptide of the correct mass appeared in the presence of active kinase, concomitant with depletion of the confidently identified unmodified peptide, confirming that the peptide was phosphorylated by recombinant PfCDPK4 (Supplementary Data S4). (b) The suitability of a synthetic peptide as a PfCDPK4 substrate was quantified as the percent of the peptide that was phosphorylated over the duration of the *in vitro* reaction with calcium-activated kinase relative to the same reaction carried out with kinase that was inactivated with a calcium chelator. The p-value is given for triplicate analyses carried out with active and inactive kinase. (c) *In vivo* evidence for hypophosphorylation of the same phopshosite is shown. WT:KO is the ratio of the phosphopeptide ion peak area observed in activated wild type vs *Pfcdpk4^−^* gametocytes. “WT only” indicates the ion was not observed in *Pfcdpk4^−^* gametocytes. Where applicable, the p-value is given along with the false discovery rate (FDR) associated with multiple hypothesis testing as assessed by the Benjamini-Hochberg method (Supplementary Data S1).

## DISCUSSION

Gametocytes are sexual stages of the malaria parasite that must be taken up by mosquitoes and undergo fertilization for the completion of the parasite’s lifecycle. Upon encountering cellular triggers in the mosquito midgut, gametocytes egress out of erythrocytes and fuse to form zygotes. Our study demonstrates that PfCDPK4 is essential for male gametogenesis.

CDPKs are key mediators of Ca^2+^ signaling in the malaria parasite and function at various life cycle stages (20). The unique architecture of *Plasmodium* CDPKs that differentiates them from mammalian calmodulin-dependent protein kinases makes CDPKs attractive drug targets (10). PfCDPK1 and PfCDPK5 are involved in invasion and egress processes, respectively, during asexual blood stage infection (21, 22). PfCDPK1 and PfCDPK2 are involved in gametogenesis and are critical for establishing infection of the mosquito vector (23, 24). PfCDPK7 is an effector of phosphatidylinositol phosphate (PIP) signaling and interacts with PI(4,5)P2 and is regulating for intra-erythrocytic parasite development (25). In the rodent malaria parasite Pb, PbCDPK3 regulates ookinete motility for the invasion of the mosquito midgut (26). PbCDPK6 regulates the sporozoite’s switch from migratory to invasive upon sensing high levels of hepatocyte-specific sulfate proteoglycans (HSPGs) and functions in liver stage infection (27). *Pb cdpk4* gene disruption leads to defects in male gametogenesis and mosquito transmission (8). Other kinases such as PfPKG and PfMAP2 play a role in gametogenesis (28, 29) indicating the importance of phosphor-signaling events in sexual development of the parasite.

In the present study, we show that PfCDPK4 is expressed throughout asexual blood stage development, gametocyte development, in activated gametocytes, and sporozoite stages of the parasite. The protein primarily displays membrane localization but also shows some cytoplasmic staining in asexual and sexual stages. Myristoylation and palmitoylation at the N-termini of CDPKs have been shown to be critical for membrane targeting in plants (30) and *Plasmodium* spp. (31). Similarly, N-terminal myristoylation on glycine at position 2 in PfCDPK4 may be responsible for its membrane localization in various parasite stages. A recent study in Pb observed expression of CDPK1, CDPK4 and CDPK5 in sporozoites and described a role for PbCDPK4 in sporozoite motility, substrate attachment, and subsequent infection of hepatocytes (32). Since PfCDPK4 also displays membrane localization in sporozoites, it will be interesting to explore its role in Pf sporozoite motility and hepatocyte infection using a conditional knockout of *PfCDPK4*.

Previous studies with a conditional protein depletion system using a destabilizing domain (DD) have shown that PfCDPK4-HA-DD parasites do not show any growth defect in asexual blood stages (22). However, since the level of protein knock down using this system is not complete and residual levels of kinase may be able to perform its cellular and molecular functions, this study left the possibility of an important function for PfCDPK4 in asexual blood stages. Other studies on PfCDPK4 utilized the bumped kinase inhibitors, BKI-1 and 1294, which blocked exflagellation of gametocytes and oocyst formation in the mosquito midgut (11, 12). However, since these inhibitors can also have off-target effects on other parasite kinases, the results cannot be unequivocally interpreted. We thus sought to study the role of *PfCDPK4* by deleting the gene entirely using CRIPSR/Cas9 based gene editing. This work revealed that, although PfCDPK4 is expressed in asexual blood stage and throughout gametocyte development, it is not required for asexual blood stage replication or gametocyte development. It is possible that other CDPKs in the parasite may have a compensatory or redundant roles and activity in these stages. We however demonstrate that CDPK4 has a critical role in male gamete production, specifically the formation of flagellated microgametes that are able to individuate and egress from the activated microgametocyte. Activated male *Pf cdpk4^−^* gametocytes showed the typical morphological changes that lead to the formation of a spheroid cell upon activation, but no exflagella formation was observed in the parasites. In contrast, we observed no discernible defect in *Pf cdpk4^−^* macrogamete formation.

The generation of *Pf cdpk4^−^* parasites afforded us the unique opportunity to explore the phosphosignaling cascade initiated by activation of PfCDPK4. We used quantitative proteomics to identify phosphorylation events in gametocytes that were activated by XA, a compound endogenous to the mosquito midgut that activates CDPKs by initiating a release of intracellular calcium (6, 7). We identified many phosphopeptides that decreased in intensity or were undetectable in activated *Pf cdpk4^−^* parasites relative to WT gametocytes. We hypothesized that these hypophosphorylated proteins would include direct substrates of PfCDPK4 and indirect targets within the signaling cascade. In our experiment, gametocytes were incubated with XA for 10 minutes prior to snap-freezing and processing for proteomics. Male gametocytes undergo a drastic transformation within that time frame, including three rounds of genome replication and assembly of flagella. By contrast, *Pf cdpk4^−^* parasites failed to develop male gametes. It has been shown that the release of Ca^2+^ happens within seconds of activation by XA (8), and work in Pb has shown that PbCDPK4 has multiple windows of activity over that time frame, playing roles in both DNA replication and axoneme motility (19). It is therefore likely that deleting *Pf cdpk4* perturbs a phosphosignaling network beyond the immediate activity of the one kinase, and that hypophosphorylated proteins observed in *Pf cdpk4^−^* parasites may represent not only substrates of PfCDPK4, but also substrates of kinases downstream in the signaling cascade initiated by activation of PfCDPK4. This hypothesis is supported by our gene ontology analysis of proteins that were hypophosphorylated in the absence of PfCDPK4. These proteins were annotated with molecular functions including protein kinase activity, replication, transcription, mRNA processing, translation, and motility; i.e., the processes required for male gametogenesis (Figure 7).

**Figure 7.**
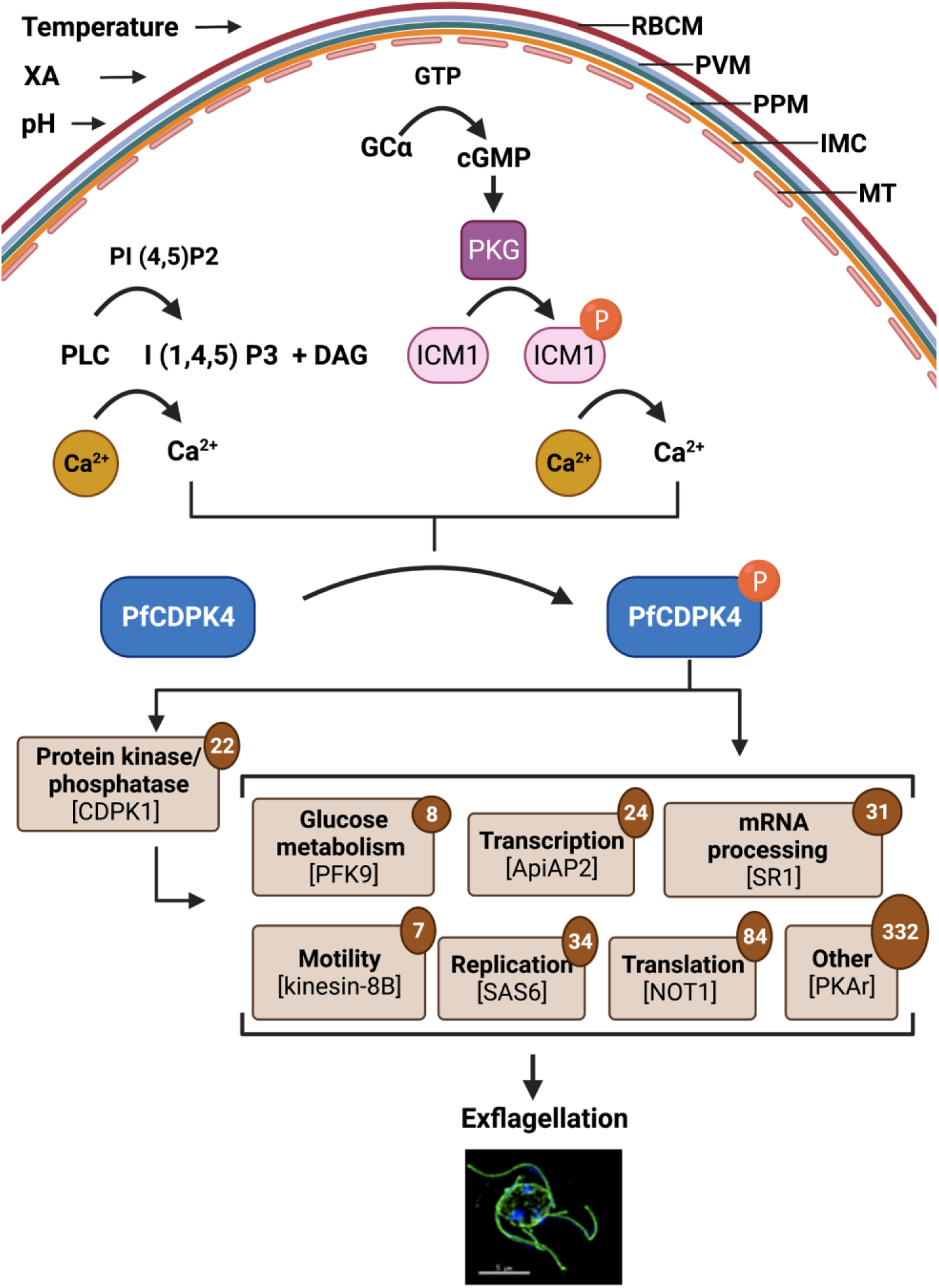
A model for the phosphosignaling network regulated by PfCDPK4. PLC-mediated and PKG-ICM1 mediated Ca^2+^ release from Ca^2+^ stores results in the activation of PfCDPK4, which may trigger phosphorylation of potential substrates belonging to protein kinase/ phosphatases, glucose metabolism, transcription, mRNA processing, motility, replication, translation and other parasite proteins. These phosphorylation events may contribute to the process of exflagellation. Numbers within circles indicated total number of proteins in that category from among the proteins we identified as hypophosphorylated in activated *Pfcdpk4^−^* gametocytes relative to wild type. Annotated and predicted protein function was based on gene ontology terms. RBCM-Red blood cell membrane, PVM-parasitophorous vacuolar membrane, PPM-parasite plasma membrane, IMC-inner membrane complex, MT-microtubular network.

In addition to elucidating the phosphosignaling network downstream of PfCDPK4 kinase activity, we also sought to identify proteins that are direct substrates of PfCDPK4. To this end, we selected several phosphosites for validation by *in vitro* kinase assays with recombinant PfCDPK4. We reasoned that if a phosphosite was hypophosphorylated in activated *Pfcdpk4^−^* gametocytes, and if that same residue was phosphorylated by recombinant PfCDPK4 *in vitro*, it is highly likely that the protein is a bona fide substrate of PfCDPK4. Our selection of targets for testing was guided by our experimental evidence as well as previously published work in other systems and life cycle stages. For example, we have here identified an uncharacterized protein, PF3D7_0417600, as a putative substrate of PfCDPK4. The protein was an especially interesting find because it is likely gametocyte-specific (it has only been detected in gametocytes in the compendium of proteomics experiments compiled at PlasmoDB.org (33)). The phosphosite S^23^ was observed *in vivo* on a phosphopeptide that was highly abundant in activated WT gametocytes, but that was significantly less abundant in *Pfcdpk4^−^* gametocytes (Table 1; Supplementary Data S1). This residue is found in a Simple 1 motif, making it a good candidate for a CDPK substrate. Our *in vitro* assay showed that the peptide was readily phosphorylated by recombinant PfCDPK4 *in vitro* (Table 1, Supplementary data S3). This protein has no predicted function or homology domains, making it a compelling target for future studies to determine its role in gametogenesis.

Using this strategy of hypothesis generation by phosphoproteomics followed by validation with *in vitro* assays, we provide evidence that substrates of PfCDPK4 include proteins previously annotated as putative SOCs by work in the rodent malaria model Pb (19), namely PfSOC3, PfSOC7, and PfPFK9. Similar to our novel SOC PF3D7_0417600, PfSOC3 has only been detected in gametocyte stages. PbSOC3 has been shown to be required for exflagellation (19). PfSOC3 may therefore be a key player in the regulation of exflagellation by PfCDPK4. PfSOC7 (also known as ribonucleoside-diphosphate reductase large subunit) plays a key role in DNA synthesis (34). Chemical inhibition of PbCDPK4 soon after activation of Pb gametocytes was shown to inhibit the genome replication that is required for the three rounds of cell division characteristic of male gametogenesis (19). It is therefore reasonable to hypothesize that this cell division event in Pf gametes may be regulated in part by phosphorylation of PfSCO7 by PfCDPK4. The enzyme 6-phospho-1-fructokinase (PFK) plays a key role in glycolysis. As *Plasmodium* male gametes lack mitochondria, glycolysis is the likely exclusive source of energy for flagellar motion (35). Since PfPFK9 possess all the catalytic features appropriate for PFK activity, it is possible that PfCDPK4-mediated phosphorylation of PfPFK9 is required for the exflagellation process in *Plasmodium* gametocytes. These results demonstrate the usefulness of the approach we have employed here for identifying novel CDPK4 substrates. We have only tested a small number of candidates within the scope of this study, but the list of hypophosphorylated phosphosites we have identified represents a wealth of testable hypotheses for future studies, including many uncharacterized proteins. Other potential targets for validation include additional putative SOC proteins that were hypophosphorylated in our *Pfcdpk4^−^* gametocytes, such as kinesin 8-B (PF3D7_0111000) and SOC9 (PF3D7_0515600), recently characterized as gamete egress protein (GEP) (36). Kinesin 8-B is a motor protein required for axoneme assembly and flagellum formation; deletion of kinesin 8-B in Pb completely blocked transmission (37). PbGEP has roles in both male and female gametocyte fertility, and deletion of PbGEP prevented transmission by causing male gametes to be immotile.

Finally, another compelling observation that arose from our phosphoproteomic analysis is evidence for crosstalk among CDPKs in activated gametocytes. PfCDPK1 is another kinase from the parasite CDPK family that is critical for exflagellation (23). We identified 13 PfCDPK1 phosphosites in our phosphoprotoemic analysis of activated gametocytes, including hypophosphorylation of four sites in *Pf cdpk4^−^* parasites: S^64^, S^204^, S^220^, which *in vitro* studies have indicated are autophosphorylation sites; and T^231^, mutation of which has been shown to lead to reduced kinase activity *in vitro* (16). In contrast to other putative SOCs discussed above, the magnitude of hypophosphorylation of these sites was more subtle, on the order of 1.5− to 6.5-fold lower in *Pf cdpk4^−^* parasites compared to WT parasites. However, our *in vitro* assay showed that a peptide bearing PfCDPK1 S^64^, which is found in a Simple 1 motif, was readily phosphorylated by recombinant PfCDPK4. These findings indicate the possibility of crosstalk between PfCDPK4 and PfCDPK1 as the two kinases mediate gametogenesis in the parasite.

The work we describe here highlights the intricate phosphosignaling mechanisms in Pf gametocytes that may regulate the function of key parasite proteins in male gametogenesis. Given the role of PfCDPK4 in male gametogenesis and transmission of the parasite to the mosquito vector, this protein and its substrates are prime targets for transmission-blocking strategies to aid malaria elimination efforts.

## MATERIALS AND METHODS

### Reagents and primary antibodies

All molecular biology reagents and oligonucleotides were purchased from Millipore Sigma (USA) unless otherwise stated. The following primary antibodies and antisera and dilutions were used: mouse anti-tubulin antibody (1:250, Millipore Sigma, cat# T5168); mouse anti-CSP (clone 2A10, 1:250), mouse anti-Pfg377 (1:250, kindly gifted by Professor Pietro Alano at Istituto Superiore di Sanità, Italy). Anti-PfUIS4 antisera is described in (38). The generation of polyclonal rabbit antisera against PfCDPK4 (1:100) is described below.

### *P. falciparum* culture and transfection

*P. falciparum* NF54 and *Pfcdpk4^−^* parasites were cultured as asexual blood stages according to standard procedures and received complete RPMI media supplemented either with 0.5% AlbuMAX^TM^ (Thermo Scientific) medium or 10% (v/v) human serum changes every 24 hrs. Gametocyte cultures were set up in 6 well plates using O^+^ human RBCs (Valley Biomedical, VA, US) and O^+^ human serum (Interstate Blood Bank, TN, US) with a final volume of 5 mL at 1% starting parasitemia and 5% hematocrit. Cultures were kept at 37°C and supplemented with gas containing 5% O_2_/5% CO_2_/90% N_2_.

Oligonucleotide primers used for the creation and analysis of *Pfcdpk4^−^* parasites are detailed in Supplementary Data S5. Deletion of *PfCDPK4* (PlasmoDB identifier Gene – PF3D7_0717500) was achieved based on the previously reported CRISPR/Cas9 strategy using the pFC plasmid (39). The *PfCDPK4* locus was deleted using double crossover homologous recombination. Complementary regions of *PfCDPK4* upstream and downstream of the open reading frame were ligated into plasmid pFCL3 (generated by modification of the pYC plasmid) as was the 20-nucleotide guide RNA sequence resulting in the creation of plasmids pFCL3_CDPK4_KO. 100 μg of this plasmid DNA was transfected into the Pf NF54 ring stage parasites vie electroporation at 310 V and 950 μF by using a Bio-Rad Gene Pulser II (Bio-Rad Laboratories, Hercules, CA) and selected using 8 nM WR99210 (kindly gifted by Jacobus Pharmaceuticals). Gene deletion was confirmed by genotyping PCR (see Fig. 3C). Two individual clones for *Pfcdpk4^−^* (clone 1C2 and 1D2) were used for phenotypic analysis.

### Generation of antisera

Amino acids 329–343 of the J-domain of PfCDPK4 were conjugated to carrier protein Keyhole limpet hemocyanin (KLH) (KMMTSKDNLNIDIPS -KLH) was used for immunization of rabbits by YenZym Antibodies, LLC (CA, US). Antibody titers were measured by ELISA by YenZym Antibodies, LLC.

### Measurement of asexual blood stage growth and gametocyte development

To compare asexual blood stage replication and growth between the NF54 WT and *Pfcdpk4^−^* parasites, synchronized parasites were set up at an initial parasitemia of 1% at ring stages and cultured in 6-well plates as described above. Parasites were removed at 48 and 96 hrs. for preparation of Giemsa-stained thin blood smears and parasitemia was scored per 1000 erythrocytes. To compare gametocyte formation between WT NF54 and *Pfcdpk4^−^*, gametocytes were cultured as described elsewhere (40). Parasites were removed on day 15 of *in vitro* culture for preparation of Giemsa-stained thin blood smears and gametocytemia was scored per 1000 erythrocytes.

### Indirect immunofluorescence

For IFAs of gametocytes and exflagellating gametes, thin smears were prepared on Teflon coated slides and fixed with 4% paraformaldehyde/0.0025% glutaraldehyde solution for 30 min. Slides were kept in a humidity chamber for each step. Fixed parasites were washed twice with PBS and permeabilized using 0.1% Triton X-100/PBS solution for 10 min. Parasites were washed with PBS and blocked with 3%BSA/PBS for 45 min. Primary antisera in 3% BSA/PBS was added to the parasites and slides were incubated at 4°C. Antigens were visualized using anti-species antibodies. Images were obtained using a 100× 1.4 NA objective 90 (Olympus) on a Delta Vision Elite High-Resolution Microscope (GE Healthcare Life Sciences).

For IFAs of sporozoites, sporozoites were resuspended in Schneider’s Drosophila Medium (Thermo Scientific) and ~20,000 parasites were spotted per well on poly-lysine coated slides (Cat# 48382-117), and air dried overnight. Parasites were fixed in 4% paraformaldehyde for 15 min and permeabilized in 0.5% Triton X-100 for 5 min. Sporozoites were blocked in 3% BSA/PBS for 1 hr. prior to incubation with antibodies. Primary antisera in 3% BSA/PBS was added to the parasites and slides were incubated at 4°C. Antigens were visualized using anti-species antibodies (Alexa Fluor, Thermo Scientific). Images were obtained using a 100× 1.4 NA objective 90 (Olympus) on a Delta Vision Elite High-Resolution Microscope (GE Healthcare Life Sciences).

### Statistical analysis of molecular parasitology data

All data related to phenotyping assays are expressed as mean ± SD. Statistical differences were deemed significant using p-values from an unpaired, two-tailed Student’s t-test. Values of p < 0.05 were considered statistically significant. Significances were calculated using GraphPad Prism 8 and are represented in the Figures as follows: ns, not significant, p > 0.05; *p < 0.05; **p < 0.01; ***p < 0.001.

### Preparation of WT and *Pf cdpk4^−^* gametocytes for LC-MS/MS

WT NF54 and *Pf cdpk4^−^* gametocytes were cultured as described above. Stage V gametocytes were activated for 10 mins by addition of 100 μM xanthurenic acid (XA) (Millipore SIGMA) and harvested using saponin lysis. Parasite pellets were washed with ice-cold PBS to remove RBC proteins and stored at −80°C until further analysis. The four activated gametocyte samples (two biological replicates each of WT and *Pf cdpk4^−^*) were processed in parallel. Unless indicated otherwise, all chemical reagents used for proteomic sample preparation were sourced from Millipore Sigma (USA), and water and acetonitrile (ACN) were LC-MS grade from Honeywell Burdick & Jackson (USA). To each pellet was added an equal volume of 2× lysis buffer containing 4% sodium dodecyl sulfate (SDS), 100 mM ammonium bicarobonate (ABC), 100 mM tris-carboxyethylphosphine (TCEP; Thermo Fisher Scientific #77720), and 2× Halt Protease and Phosphatase Inhibitor Cocktail, EDTA-free (Thermo Fisher Scientific # 78441). The pellets were incubated for 10 min in a 95°C water bath with intermittent vortexing. Iodoacetamide was added to a final concentration of 70 mM and the samples were incubated in darkness at room temperature for 30 min with vortexing. Insoluble debris was pelleted by centrifuging 5 min at 20,000×g and supernatants were transferred to clean tubes. Protein was precipitated by the Wessel-Flugge method (41) and resolubilized in 8M urea/50 mM ABC. Protein concentration was determined using the Pierce BCA assay (Thermo Fisher Scientific #23225), and four 500 μg aliquots of each sample were taken for digestion. Each aliquot was diluted eight-fold with 50 mM ABC and 10 μg of sequencing grade modified trypsin (Promega #V5113) was added. The samples were incubated overnight on a thermomixer at 37°C, desalted on Pierce peptide desalting spin columns (Thermo Fisher Scientific # 89851), and dried in a vacuum concentrator.

### Phosphopeptide enrichment

Phosphopeptides were enriched automatically using magnetic immobilized metal affinity chromatography (IMAC) beads (ReSyn biosciences, South Africa) using a Qiagen BioSprint 96 (Qiagen, Germany) magnetic particle handler. For each sample, four aliquots of peptides from 500 μg of digested protein were processed in parallel, two with Ti-IMAC HP beads and two with Zr-IMAC HP beads, and the resulting phosphopeptides were combined. Briefly, the manufacturer’s protocol was followed: desalted and dried peptides from 500 μg of digested protein were resuspended in load buffer (1.0 M glycolic acid/80% ACN/5% TFA for Ti-IMAC HP beads, 0.1 M glycolic acid/80% ACN/5% trifluoroacetic acid (TFA; Optima LC/MS grade, Fisher Chemical, USA) for Zr-IMAC HP beads) and incubated with beads. Beads were washed in load buffer, then 80% ACN/1% TFA, then 10% ACN/0.2% TFA, then eluted with 2% v/v ammonium hydroxide. The eluted peptide solution was acidified with TFA, desalted on Pierce peptide desalting spin columns (Thermo Fisher Scientific # 89851), and dried in a vacuum concentrator.

### LC-MS/MS of phosphopeptides

Purified and desalted phosphopeptides were resuspended in 80 μL of 0.1% TFA. Eight μL was injected for each LC-MS/MS analysis. Two replicate liquid chromatography-tandem mass spectrometry (LC-MS/MS) analyses were performed for each of the four samples. LC was performed with an EASY-nLC 1000 (Thermo Fisher Scietific, USA) using a vented trap set-up. The trap column was a PepMap 100 C18 (Thermo Fisher Scientific #164535) with 75 μm i.d. and a 2 cm bed of 3μm 100 Å C18. The analytical column was an EASY-Spray column (ThermoFisher Scientific #ES803revA) with 75 μm i.d. and a 50 cm bed of 2μm 100 Å C18 operated at 25°C. The LC mobile phases consisted of buffer A (0.1 % v/v formic acid in water) and buffer B (0.1 % v/v formic acid in ACN). The separation gradient, operated at 300 nL/min, was 5 % B to 35 % B over 4 h. MS/MS was performed with a Thermo Fisher Scientific Orbitrap Fusion Lumos using data-dependent acquisition (DDA). The following acquisition settings were used: MS1 scan from 375-1550 m/z at 120,000 resolution with an AGC target of 3×10^6^ ions and a max fill time of 20 ms; MS2 at 30,000 resolution with an AGC target of 1×10^6^ ions and a max fill time of 60 ms; 15 data dependent scans with a 30 sec dynamic exclusion window; select only precursors with charge +2, +3, +4, or +5 with monoisotopic precursor selection enabled; HCD fragmentation at 28% normalized collision energy, 1.6 m/z isolation window with no offset.

### Data processing for phosphopeptide identification and quantification

Complete data analysis parameters are given in Supplementary Data S6. Briefly, mass spectrometry output files were converted to mzML format with msConvert version 3.0.19106 (Proteowizard (42)) and searched with Comet version 2020.01 rev. 3 (43) against a database comprising *P. falciparum* 3D7 (44) (PlasmoDB v.51; www.plasmodb.org (33)) appended with the UniRef90 human proteome (UniProt (45)) and the common Repository of Adventitious Proteins (v.2012.01.01, The Global Proteome Machine, www.thegpm.org/cRAP). Decoy proteins with the residues between tryptic residues randomly shuffled were generated using a tool included in the TPP and interleaved among the real entries. The search parameters included a static modification of +57.02 Da at Cys for formation of S-carboxamidomethyl-Cys by iodoacetamide and potential modifications of +15.99 Da at Met for oxidation, +79.97 at Ser/Thr/Tyr for phosphorylation, and +42.01 for acetylation at the N-terminus of the protein, either at N-terminal Met or the N-terminal residue after cleavage of N-terminal Met. Search results were analyzed with TPP version 6.0.0 Noctilucent (46). Peptide spectrum matches (PSMs) were scored with iProphet (47), and only PSMs with probabilities >0.9 were taken for further analysis. Phopsphosite localization confidence was scored using PTMProphet (48), and a minimum localization probability of 0.7 for all phosphosites was required for the peptide to be taken for quantification. Phosphopeptides were quantified as individual ions, i.e., distinct positional isomers with distinct charges were quantified separately. Phosphopeptide ion quantification was performed with software developed in-house. Precursor extracted ion chromatograms (XICs) were computed for each target peptide observed from each corresponding mzML mass spectrum data file using a two-pass method. In the first pass XICs were computed only for the target peptides from each file using the 10-ppm mass tolerance and wide (5 min) retention time tolerance from the spectral identification. The most abundant XICs found in all MS runs at 5-min intervals were used to perform chromatographic alignment across runs by computing a linear retention time correction factor for each MS run that minimized the variance in retention time of these anchor peptides. This retention time alignment allowed for match-between-run (49) analysis from the master list of all peptide sequences observed in all runs in a second pass of XIC extraction. In the second pass, XICs for all target peptides, regardless of which MS run they originated from were extracted with a 10-ppm mass tolerance and within 1 min of the average retention time among those samples where the peptide was observed.

### Data processing for determining differential phosphorylation

Only Pf phosphopeptides were taken for further consideration. Phosphopeptide ion peak areas were normalized to account for run-to-run variation using the 1881 phosphopeptide ions quantified in all eight LC-MS/MS runs. The sum of these peak areas was taken for each of the eight runs, and the run with the highest sum was used as reference run to serve as the basis for normalization. A correction factor for each of the other seven runs was calculated as follows: for each peptide ion, the log of the ratio of the peak area relative to the peak area for the same ion in the reference run was calculated, and the distribution of all log ratios was fitted to a Gaussian curve. The mean of this distribution was taken as a correction factor which was then applied to all peak areas in the respective runs. For phosphopeptide ions quantified in both WT and *Pf cdpk4^−^* gametocytes, two different approaches were used to assess significance. First, the ratio of the mean peak areas for WT and *Pf cdpk4^−^* were calculated for each ion, and the distribution of the log-transformed ratios was fitted to a Gaussian curve. The complementary error function (ERFC) was then used to calculate a p-value for each ratio and a Benjamini-Hochberg false discovery rate (FDR) was calculated. Second, for peptide ions quantified in at least two out of four runs for both WT and *Pf cdpk4^−^* gametocytes, a p-value was calculated using a two-tailed, homoschedastic t-test, and a Benjamini-Hochberg FDR was calculated. Several stringent criteria were applied in order to identify phosphosites that were significantly down-regulated in *Pf cdpk4^−^* gametocytes. Only phosphopeptide ions detected by PSMs in both WT biological replicates were considered. Further, peptide ions detected in WT gametocytes were ranked by mean peak area, and those in the bottom quartile of abundance were not taken for further consideration. The remaining ions were required to meet one of the three following criteria: 1) where a t-test could be performed, the FDR of the resulting p-value was less than 10%; 2) where a ratio could be calculated but a t-test could not be performed due to insufficient data points, the FDR from the ERFC p-value was less than 10%; 3) the peptide ion was only quantified in WT and not in *Pf cdpk4^−^* gametocytes.

### Gene ontology analysis

All proteins associated with phosphosites identified as significantly hypophosphorylated were subjected to gene ontology (GO) analysis (50) using the tool available through PlasmoDB.org. The following settings were used: computed and curated evidence allowed; use GO Slim terms; P-value cutoff = 0.05.

### Motif analysis

All phosphosites identified as significantly hypophosphorylated (excluding CDPK4) were analyzed for enrichment of amino acid sequence motifs using MoMo(51) as implemented in The MEME Suite (52) (meme-suite.org). The MoDL algorithm was used with default settings.

### *In vitro* kinase assays

The recombinant maltose-binding protein (MBP) tag-fused *Pf*CDPK4 protein used for *in vitro* assays has been described previously (11). Protein kinase assays were performed using synthetic peptides corresponding to various putative PfCDPK4 substrates (Supplementary Data S3). Synthetic peptides (purity ≥90%) were ordered from Biomatik (Ontario, Canada). Syntide 2 (Millipore Sigma #SCP0250) was used as a positive control. Peptides were reconstituted in water or DMSO as required for solubility and pooled into a stock solution with each peptide at 50 μM. The resulting concentration of DMSO in this stock solution was 13.5% (v/v). Kinase reaction buffer prepared at 2× concentration contained 100 mM Tris pH 7.5, 20 mM MgCl_2_, 2mM DTT (Thermo Fisher Scientific), 4 mM ATP (Thermo Fisher Scientific #R1441), and either 4 mM CaCl_2_ (to activate the kinase) or 4 mM EGTA (to keep the kinase inactive for the control). Triplicate reactions with either the CaCl_2_ buffer or the EGTA (Fluka, Germany) buffer were performed in parallel. Recombinant MBP-tagged PfCDPK4 (4.14 μL of a 4.84μM stock in 50% glycerol) and synthetic peptides (10 μL of the pooled 50 μM peptide stock) were diluted to 50 μL with water, and the reaction was initiated by adding 50 μL of kinase buffer. The reaction was incubated on a thermomixer at 30°C for 1 h. The reaction was stopped by taking 10 μL of the reaction solution and adding it to 10 μL of 1% TFA/10% ACN on ice. The acidified solution was then desalted on a C18 spin column (Pierce) and dried in a vacuum concentrator.

### LC-MS/MS analysis of *in vitro* kinase assays

Desalted peptides from the kinase assays were resuspended in 50 μL of 0.1% TFA. Three μL were injected for each LC-MS/MS analysis. Liquid chromatography was performed with an EASY-nLC 1000 (Thermo Fisher Scientific) using a vented trap set-up. The trap column was a PepMap 100 C18 (Thermo Fisher Scientific #164535) with 75 μm i.d. and a 2 cm bed of 3μm 100 Å C18. The analytical column was an EASY-Spray column (Thermo Fisher Scientific #ES804revA) with 75 μm i.d. and a 15 cm bed of 2μm 100 Å C18 operated at 25°C. The LC mobile phases consisted of buffer A (0.1 % v/v formic acid in water) and buffer B (0.1 % v/v formic acid in ACN). The separation gradient, operated at 400 nL/min, was 5 % B to 35 % B over 30 min. MS/MS was performed with a Thermo Fisher Scientific Q-Exactive HF Orbitrap using DDA. An inclusion list of 255 m/z values was employed to prioritize selection of precursors matching phosphopeptide isoforms with any number of modified Ser or Thr residues observed at charge +2, +3, or +4. The following acquisition settings were used: MS1 scan from 375-1075 m/z at 60,000 resolution with an AGC target of 3×10^6^ ions and a max fill time of 20 ms; MS2 at 30,000 resolution with an AGC target of 1×10^6^ ions and a max fill time of 100 ms; DDA loop count of 25 with no dynamic exclusion; select only precursors with charge +2, +3, or +4, exclude isotopes and peptide match enabled; use inclusion list, select other precursors if idle; HCD fragmentation at 27% normalized collision energy, 1.8 m/z isolation window with no offset.

### Data processing for *in vitro* kinase assays

Complete data analysis parameters are given in Supplementary Data S6. Briefly, mass spectrometry output files were converted to mzML format with msConvert version 3.0.19106 (Proteowizard (42)) and searched with Comet version 2020.01 rev. 3(43) against a database containing the sequences of the synthetic peptides, allowing for variable modification of of +79.97 at Ser/Thr/Tyr and +15.99 at Met. Search results were analyzed with the Trans Proteomic Pipeline (TPP) version 6.0.0 Noctilucent (46). Peptide peak heights were quantified with the XPRESS tool in the TPP using the label-free option. PSMs with PeptideProphet probabilities >0.9 were used to identify the peptide peaks. Phosphosites were considered positively localized if the PTMProphet localization score was >0.7. Missing peptide peak height values (e.g. for low-abundance unmodified peptides depleted by conversion to phosphopeptide) were manually determined from the extracted ion chromatogram at the observed retention time. Peptide peak heights in the presence of active kinase (CaCl_2_ kinase buffer) or inactive kinase (EGTA kinase buffer) were taken as the mean of the technical triplicates. The percentage of the peptide that was converted to phosphopeptide was quantified as the ratio of unphosphorylated peptide peak height in the presence of active kinase relative to the peak height in the presence of inactive kinase. The kinase activity for a given peptide was reported as 1 minus this ratio, i.e. the percent of the peptide that was converted to phosphopeptide. A p-value was assigned using a one-tailed homoschedastic t-test. Peptides were considered good substrates of PfCDPK4 if >70% of the peptide was converted to phosphopeptide and the p-value was <0.05.

### PfCDPK4 autophosphorylation assay

Recombinant MBP-tagged PfCDPK4 was incubated in CaCl_2_ kinase buffer as above without peptide substrates present. IAM was added to a final concentration of 20 mM and the solution was incubated with vortexing for 20 min at room temperature in darkness, after which 0.1 mg of sequencing grade modified trypsin was added. The solution was incubated with vortexing for 2 h at 37°C. ACN and TFA were added to final concentrations of 5% v/v and 0.5% v/v, respectively, and the peptides were de-salted on a C18 spin column (G-Biosciences #786-930) and dried in a vacuum concentrator.

### LC-MS/MS analysis of PfCDPK4 autophosphorylation assay

Desalted peptides from the recombinant PfCDPK4 digest were resuspended in 40 μL of 0.1% TFA. Four μL were injected for each of two replicate LC-MS/MS analyses. Liquid chromatography was performed as above. MS/MS was performed as above with the following differences: no inclusion list; MS1 scan from 375-1375 m/z; DDA loop count of 15 with dynamic exclusion for 15 s after a single observation.

### Data processing for PfCDPK4 autophosphorylation assay

Complete data analysis parameters are given in Supplementary Data S6. Raw data were converted with MSConvert, searched with Comet, and analyzed with the TPP as above. Spectra were searched against a database comprising the sequence of the recombinant MBP-tagged PfCDPK4, the UniProt *E. coli* reference proteome, and the cRAP contaminants database, allowing for variable modification of 79.99 at STY. PfCDPK4 phosphopeptides were taken for consideration if they met the following criteria: identified by two or more PSMs with PeptideProphet corresponding to a decoy-estimated FDR<1% among phosphopeptide PSM, two tryptic termini, and phosphosite localization confirmed with a PTMProphet score >0.7.

## Data availability

The mass spectrometry data generated in the manuscript has been deposited to the ProteomeXchange Consortium (http://proteomecentral.proteomexchange.org) and can be accessed via the MassIVE partner repository (https://massive.ucsd.edu/) with the following data identifiers: MSV000087569 and PXD026747 for the WT and *Pf cdpk4^−^* gametocytes data; MSV000087575 and PXD026499 for the *in vitro* kinase assay data. All other relevant data are available from the authors upon request.

## Ethical statement

The animal experiments, which involved antisera generation in rabbits, were performed at YenZym Antibodies, LLC (CA, US) and prescribed guidelines were followed.

## Supplementary Information

Supplementary Data S1-S6.

## GRANTS

NIH NIAID K25AI119229 (KES)

NIH NIAID R01AI148489 (KES)

NIH NIAID P01 AI27338 (AMV, SHIK)

R21AI123690 (KKO)

NIH NIGMS R01GM087221 (RLM)

NIH S10OD026936 (RLM)

NSF DBI-1920268 (RLM)

## Author Contributions

Sudhir Kumar SK

Meseret T. Haile MTH

Michael R. Hoopmann MRH

Linh T. Tran LTT

Samantha A. Michaels SAM

Seamus R. Morrone SRM

Kayode K Ojo KKO

Laura M. Reynolds LMR

Ulrike Kusebauch UK

Ashley M. Vaughan AMV

Robert L. Moritz RLM

Stefan H.I. Kappe SHIK

Kristian E. Swearingen KES

Conceptualization: SK KES

Methodology: SK MRH SRM KKO UK KES

Software: MRH

Formal Analysis: SK KKO SHIK KES

Investigation: SK MTH LTT SAM KKO LMR KES

Resources: KKO RLM SHIK KES

Data Curation: SK KES

Writing – Original Draft Preparation: SK KES

Writing – Review & Editing: SK MTH MRH LTT SAM SRM KKO LRM UK AMV RLM SHIK KES

Visualization: SK KES

Supervision: KKO AMV RLM SHIK KES

Project Administration: SHIK KES

Funding Acquisition: KKO AMV RLM SHIK KES

## REFERENCES

1. Brancucci NMB, Gerdt JP, Wang C, De Niz M, Philip N, Adapa SR, Zhang M, Hitz E, Niederwieser I, Boltryk SD, Laffitte MC, Clark MA, Grüring C, Ravel D, Blancke Soares A, Demas A, Bopp S, Rubio-Ruiz B, Conejo-Garcia A, Wirth DF, Gendaszewska-Darmach E, Duraisingh MT, Adams JH, Voss TS, Waters AP, Jiang RHY, Clardy J, Marti M. 2017. Lysophosphatidylcholine Regulates Sexual Stage Differentiation in the Human Malaria Parasite Plasmodium falciparum. Cell 171:1532–1544.e15.

2. Wein S, Ghezal S, Buré C, Maynadier M, Périgaud C, Vial HJ, Lefebvre-Tournier I, Wengelnik K, Cerdan R. 2018. Contribution of the precursors and interplay of the pathways in the phospholipid metabolism of the malaria parasite. J Lipid Res 59:1461–1471.

3. Aly AS, Vaughan AM, Kappe SH. 2009. Malaria parasite development in the mosquito and infection of the mammalian host. Annu Rev Microbiol 63:195–221.

4. Sinden RE, Croll NA. 1975. Cytology and kinetics of microgametogenesis and fertilization in Plasmodium yoelii nigeriensis. Parasitology 70:53–65.

5. Sinden RE. 1983. Sexual development of malarial parasites. Adv Parasitol 22:153–216.

6. Billker O, Lindo V, Panico M, Etienne AE, Paxton T, Dell A, Rogers M, Sinden RE, Morris HR. 1998. Identification of xanthurenic acid as the putative inducer of malaria development in the mosquito. Nature 392:289–92.

7. Garcia GE, Wirtz RA, Barr JR, Woolfitt A, Rosenberg R. 1998. Xanthurenic acid induces gametogenesis in Plasmodium, the malaria parasite. J Biol Chem 273:12003–5.

8. Billker O, Dechamps S, Tewari R, Wenig G, Franke-Fayard B, Brinkmann V. 2004. Calcium and a calcium-dependent protein kinase regulate gamete formation and mosquito transmission in a malaria parasite. Cell 117:503–14.

9. Balestra AC, Koussis K, Klages N, Howell SA, Flynn HR, Bantscheff M, Pasquarello C, Perrin AJ, Brusini L, Arboit P, Sanz O, Castaño LP, Withers-Martinez C, Hainard A, Ghidelli-Disse S, Snijders AP, Baker DA, Blackman MJ, Brochet M. 2021. Ca(2+) signals critical for egress and gametogenesis in malaria parasites depend on a multipass membrane protein that interacts with PKG. Sci Adv 7.

10. Billker O, Lourido S, Sibley LD. 2009. Calcium-dependent signaling and kinases in apicomplexan parasites. Cell Host Microbe 5:612–22.

11. Ojo KK, Pfander C, Mueller NR, Burstroem C, Larson ET, Bryan CM, Fox AM, Reid MC, Johnson SM, Murphy RC, Kennedy M, Mann H, Leibly DJ, Hewitt SN, Verlinde CL, Kappe S, Merritt EA, Maly DJ, Billker O, Van Voorhis WC. 2012. Transmission of malaria to mosquitoes blocked by bumped kinase inhibitors. J Clin Invest 122:2301–5.

12. Ojo KK, Eastman RT, Vidadala R, Zhang Z, Rivas KL, Choi R, Lutz JD, Reid MC, Fox AM, Hulverson MA, Kennedy M, Isoherranen N, Kim LM, Comess KM, Kempf DJ, Verlinde CL, Su XZ, Kappe SH, Maly DJ, Fan E, Van Voorhis WC. 2014. A specific inhibitor of PfCDPK4 blocks malaria transmission: chemical-genetic validation. J Infect Dis 209:275–84.

13. Govindasamy K, Jebiwott S, Jaijyan DK, Davidow A, Ojo KK, Van Voorhis WC, Brochet M, Billker O, Bhanot P. 2016. Invasion of hepatocytes by Plasmodium sporozoites requires cGMP-dependent protein kinase and calcium dependent protein kinase 4. Mol Microbiol 102:349–363.

14. Hegeman AD, Rodriguez M, Han BW, Uno Y, Phillips GN, Jr., Hrabak EM, Cushman JC, Harper JF, Harmon AC, Sussman MR. 2006. A phyloproteomic characterization of in vitro autophosphorylation in calcium-dependent protein kinases. Proteomics 6:3649–64.

15. Harper JF, Harmon A. 2005. Plants, symbiosis and parasites: a calcium signalling connection. Nat Rev Mol Cell Biol 6:555–66.

16. Ahmed A, Gaadhe K, Sharma GP, Kumar N, Neculai M, Hui R, Mohanty D, Sharma P. 2012. Novel insights into the regulation of malarial calcium-dependent protein kinase 1. Faseb j 26:3212–21.

17. Ranjan R, Ahmed A, Gourinath S, Sharma P. 2009. Dissection of mechanisms involved in the regulation of Plasmodium falciparum calcium-dependent protein kinase 4. J Biol Chem 284:15267–76.

18. Treeck M, Sanders JL, Elias JE, Boothroyd JC. 2011. The phosphoproteomes of Plasmodium falciparum and Toxoplasma gondii reveal unusual adaptations within and beyond the parasites’ boundaries. Cell Host Microbe 10:410–9.

19. Fang H, Klages N, Baechler B, Hillner E, Yu L, Pardo M, Choudhary J, Brochet M. 2017. Multiple short windows of calcium-dependent protein kinase 4 activity coordinate distinct cell cycle events during Plasmodium gametogenesis. Elife 6.

20. Ghartey-Kwansah G, Yin Q, Li Z, Gumpper K, Sun Y, Yang R, Wang D, Jones O, Zhou X, Wang L, Bryant J, Ma J, Boampong JN, Xu X. 2020. Calcium-dependent Protein Kinases in Malaria Parasite Development and Infection. Cell Transplant 29:963689719884888.

21. Kumar S, Kumar M, Ekka R, Dvorin JD, Paul AS, Madugundu AK, Gilberger T, Gowda H, Duraisingh MT, Keshava Prasad TS, Sharma P. 2017. PfCDPK1 mediated signaling in erythrocytic stages of Plasmodium falciparum. Nat Commun 8:63.

22. Dvorin JD, Martyn DC, Patel SD, Grimley JS, Collins CR, Hopp CS, Bright AT, Westenberger S, Winzeler E, Blackman MJ, Baker DA, Wandless TJ, Duraisingh MT. 2010. A plant-like kinase in Plasmodium falciparum regulates parasite egress from erythrocytes. Science 328:910–2.

23. Bansal A, Molina-Cruz A, Brzostowski J, Liu P, Luo Y, Gunalan K, Li Y, Ribeiro JMC, Miller LH. 2018. PfCDPK1 is critical for malaria parasite gametogenesis and mosquito infection. Proc Natl Acad Sci U S A 115:774–779.

24. Bansal A, Molina-Cruz A, Brzostowski J, Mu J, Miller LH. 2017. Plasmodium falciparum Calcium-Dependent Protein Kinase 2 Is Critical for Male Gametocyte Exflagellation but Not Essential for Asexual Proliferation. mBio 8.

25. Kumar P, Tripathi A, Ranjan R, Halbert J, Gilberger T, Doerig C, Sharma P. 2014. Regulation of Plasmodium falciparum development by calcium-dependent protein kinase 7 (PfCDPK7). J Biol Chem 289:20386–95.

26. Siden-Kiamos I, Ecker A, Nybäck S, Louis C, Sinden RE, Billker O. 2006. Plasmodium berghei calcium-dependent protein kinase 3 is required for ookinete gliding motility and mosquito midgut invasion. Mol Microbiol 60:1355–63.

27. Coppi A, Tewari R, Bishop JR, Bennett BL, Lawrence R, Esko JD, Billker O, Sinnis P. 2007. Heparan sulfate proteoglycans provide a signal to Plasmodium sporozoites to stop migrating and productively invade host cells. Cell Host Microbe 2:316–27.

28. McRobert L, Taylor CJ, Deng W, Fivelman QL, Cummings RM, Polley SD, Billker O, Baker DA. 2008. Gametogenesis in malaria parasites is mediated by the cGMP-dependent protein kinase. PLoS Biol 6:e139.

29. Hitz E, Balestra AC, Brochet M, Voss TS. 2020. PfMAP-2 is essential for male gametogenesis in the malaria parasite Plasmodium falciparum. Sci Rep 10:11930.

30. Martín ML, Busconi L. 2000. Membrane localization of a rice calcium-dependent protein kinase (CDPK) is mediated by myristoylation and palmitoylation. Plant J 24:429–35.

31. Möskes C, Burghaus PA, Wernli B, Sauder U, Dürrenberger M, Kappes B. 2004. Export of Plasmodium falciparum calcium-dependent protein kinase 1 to the parasitophorous vacuole is dependent on three N-terminal membrane anchor motifs. Mol Microbiol 54:676–91.

32. Govindasamy K, Bhanot P. 2020. Overlapping and distinct roles of CDPK family members in the pre-erythrocytic stages of the rodent malaria parasite, Plasmodium berghei. PLoS Pathog 16:e1008131.

33. Aurrecoechea C, Brestelli J, Brunk BP, Dommer J, Fischer S, Gajria B, Gao X, Gingle A, Grant G, Harb OS, Heiges M, Innamorato F, Iodice J, Kissinger JC, Kraemer E, Li W, Miller JA, Nayak V, Pennington C, Pinney DF, Roos DS, Ross C, Stoeckert CJ, Jr., Treatman C, Wang H. 2009. PlasmoDB: a functional genomic database for malaria parasites. Nucleic Acids Res 37:D539–43.

34. Chakrabarti D, Schuster SM, Chakrabarti R. 1993. Cloning and characterization of subunit genes of ribonucleotide reductase, a cell-cycle-regulated enzyme, from Plasmodium falciparum. Proc Natl Acad Sci U S A 90:12020–4.

35. Talman AM, Prieto JH, Marques S, Ubaida-Mohien C, Lawniczak M, Wass MN, Xu T, Frank R, Ecker A, Stanway RS, Krishna S, Sternberg MJ, Christophides GK, Graham DR, Dinglasan RR, Yates JR, 3rd, Sinden RE. 2014. Proteomic analysis of the Plasmodium male gamete reveals the key role for glycolysis in flagellar motility. Malar J 13:315.

36. Andreadaki M, Pace T, Grasso F, Siden-Kiamos I, Mochi S, Picci L, Bertuccini L, Ponzi M, Currà C. 2020. Plasmodium berghei Gamete Egress Protein is required for fertility of both genders. Microbiologyopen 9:e1038.

37. Zeeshan M, Ferguson DJ, Abel S, Burrrell A, Rea E, Brady D, Daniel E, Delves M, Vaughan S, Holder AA, Le Roch KG, Moores CA, Tewari R. 2019. Kinesin-8B controls basal body function and flagellum formation and is key to malaria transmission. Life Sci Alliance 2.

38. Mackellar DC, O’Neill MT, Aly AS, Sacci JB, Jr., Cowman AF, Kappe SH. 2010. Plasmodium falciparum PF10_0164 (ETRAMP10.3) is an essential parasitophorous vacuole and exported protein in blood stages. Eukaryot Cell 9:784–94.

39. Goswami D, Betz W, Locham NK, Parthiban C, Brager C, Schäfer C, Camargo N, Nguyen T, Kennedy SY, Murphy SC, Vaughan AM, Kappe SH. 2020. A replication-competent late liver stage-attenuated human malaria parasite. JCI Insight 5.

40. Tripathi AK, Mlambo G, Kanatani S, Sinnis P, Dimopoulos G. 2020. Plasmodium falciparum Gametocyte Culture and Mosquito Infection Through Artificial Membrane Feeding. J Vis Exp doi:10.3791/61426.

41. Wessel D, Flugge UI. 1984. A method for the quantitative recovery of protein in dilute solution in the presence of detergents and lipids. Anal Biochem 138:141–3.

42. Chambers MC, Maclean B, Burke R, Amodei D, Ruderman DL, Neumann S, Gatto L, Fischer B, Pratt B, Egertson J, Hoff K, Kessner D, Tasman N, Shulman N, Frewen B, Baker TA, Brusniak MY, Paulse C, Creasy D, Flashner L, Kani K, Moulding C, Seymour SL, Nuwaysir LM, Lefebvre B, Kuhlmann F, Roark J, Rainer P, Detlev S, Hemenway T, Huhmer A, Langridge J, Connolly B, Chadick T, Holly K, Eckels J, Deutsch EW, Moritz RL, Katz JE, Agus DB, MacCoss M, Tabb DL, Mallick P. 2012. A cross-platform toolkit for mass spectrometry and proteomics. Nat Biotechnol 30:918–20.

43. Eng JK, Jahan TA, Hoopmann MR. 2013. Comet: an open-source MS/MS sequence database search tool. Proteomics 13:22–4.

44. Gardner MJ, Hall N, Fung E, White O, Berriman M, Hyman RW, Carlton JM, Pain A, Nelson KE, Bowman S, Paulsen IT, James K, Eisen JA, Rutherford K, Salzberg SL, Craig A, Kyes S, Chan MS, Nene V, Shallom SJ, Suh B, Peterson J, Angiuoli S, Pertea M, Allen J, Selengut J, Haft D, Mather MW, Vaidya AB, Martin DM, Fairlamb AH, Fraunholz MJ, Roos DS, Ralph SA, McFadden GI, Cummings LM, Subramanian GM, Mungall C, Venter JC, Carucci DJ, Hoffman SL, Newbold C, Davis RW, Fraser CM, Barrell B. 2002. Genome sequence of the human malaria parasite Plasmodium falciparum. Nature 419:498–511.

45. UniProt C. 2019. UniProt: a worldwide hub of protein knowledge. Nucleic Acids Res 47:D506–D515.

46. Deutsch EW, Mendoza L, Shteynberg D, Slagel J, Sun Z, Moritz RL. 2015. Trans-Proteomic Pipeline, a standardized data processing pipeline for large-scale reproducible proteomics informatics. Proteomics Clin Appl 9:745–54.

47. Shteynberg D, Nesvizhskii AI, Moritz RL, Deutsch EW. 2013. Combining results of multiple search engines in proteomics. Mol Cell Proteomics 12:2383–93.

48. Shteynberg DD, Deutsch EW, Campbell DS, Hoopmann MR, Kusebauch U, Lee D, Mendoza L, Midha MK, Sun Z, Whetton AD, Moritz RL. 2019. PTMProphet: Fast and Accurate Mass Modification Localization for the Trans-Proteomic Pipeline. J Proteome Res 18:4262–4272.

49. Cox J, Hein MY, Luber CA, Paron I, Nagaraj N, Mann M. 2014. Accurate proteome-wide label-free quantification by delayed normalization and maximal peptide ratio extraction, termed MaxLFQ. Mol Cell Proteomics 13:2513–26.

50. Ashburner M, Ball CA, Blake JA, Botstein D, Butler H, Cherry JM, Davis AP, Dolinski K, Dwight SS, Eppig JT, Harris MA, Hill DP, Issel-Tarver L, Kasarskis A, Lewis S, Matese JC, Richardson JE, Ringwald M, Rubin GM, Sherlock G. 2000. Gene ontology: tool for the unification of biology. The Gene Ontology Consortium. Nat Genet 25:25–9.

51. Cheng A, Grant CE, Noble WS, Bailey TL. 2019. MoMo: discovery of statistically significant post-translational modification motifs. Bioinformatics 35:2774–2782.

52. Bailey TL, Boden M, Buske FA, Frith M, Grant CE, Clementi L, Ren J, Li WW, Noble WS. 2009. MEME SUITE: tools for motif discovery and searching. Nucleic Acids Res 37:W202–8.

