## Supplementary data S1-S6 for "*Plasmodium falciparum* Calcium Dependent Protein Kinase 4 is critical for male gametogenesis and transmission to the mosquito vector": Supplementary_Data_S4_in_vitro_kinase_assay_representative_analyses.pdf

**Supplementary data S4. Representative data from LC-MS/MS-based *in vitro* kinase assay**

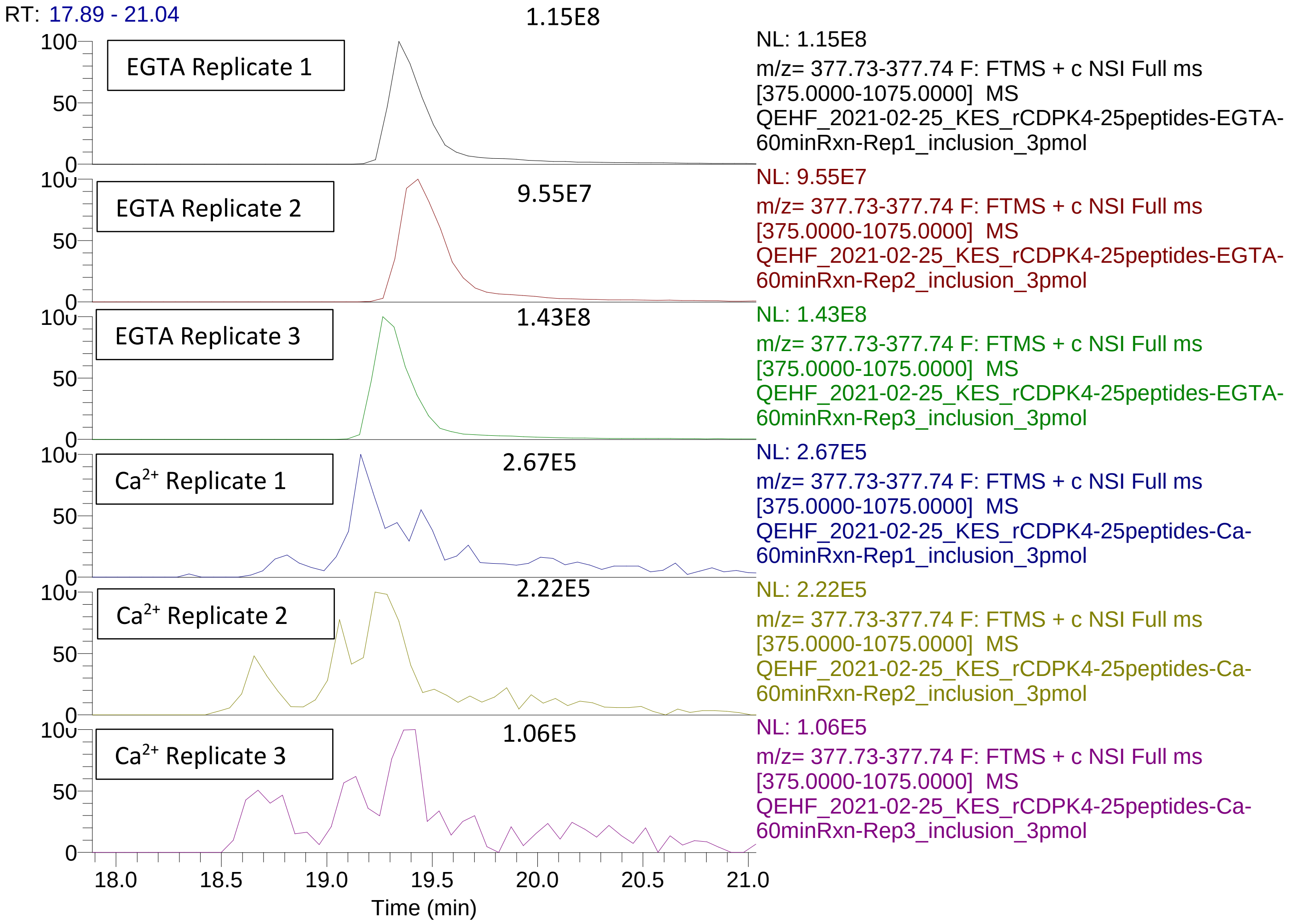


Extracted ion chromatograms for 377.74 m/z (charge +4). Peaks are labeled with peak intensity. This peptide was confidently identified in all runs as the unmodified peptide syntide 2, PLARTLSVAGLPGKK. PfCDPK4 is inactive in the presence of the calcium chelator EGTA and active in the presence of $Ca^{2+}$. In the replicates with active PfCDPK4, the peptide abundance was depleted an average of 99.8%, indicating that the peptide was phosphorylated.

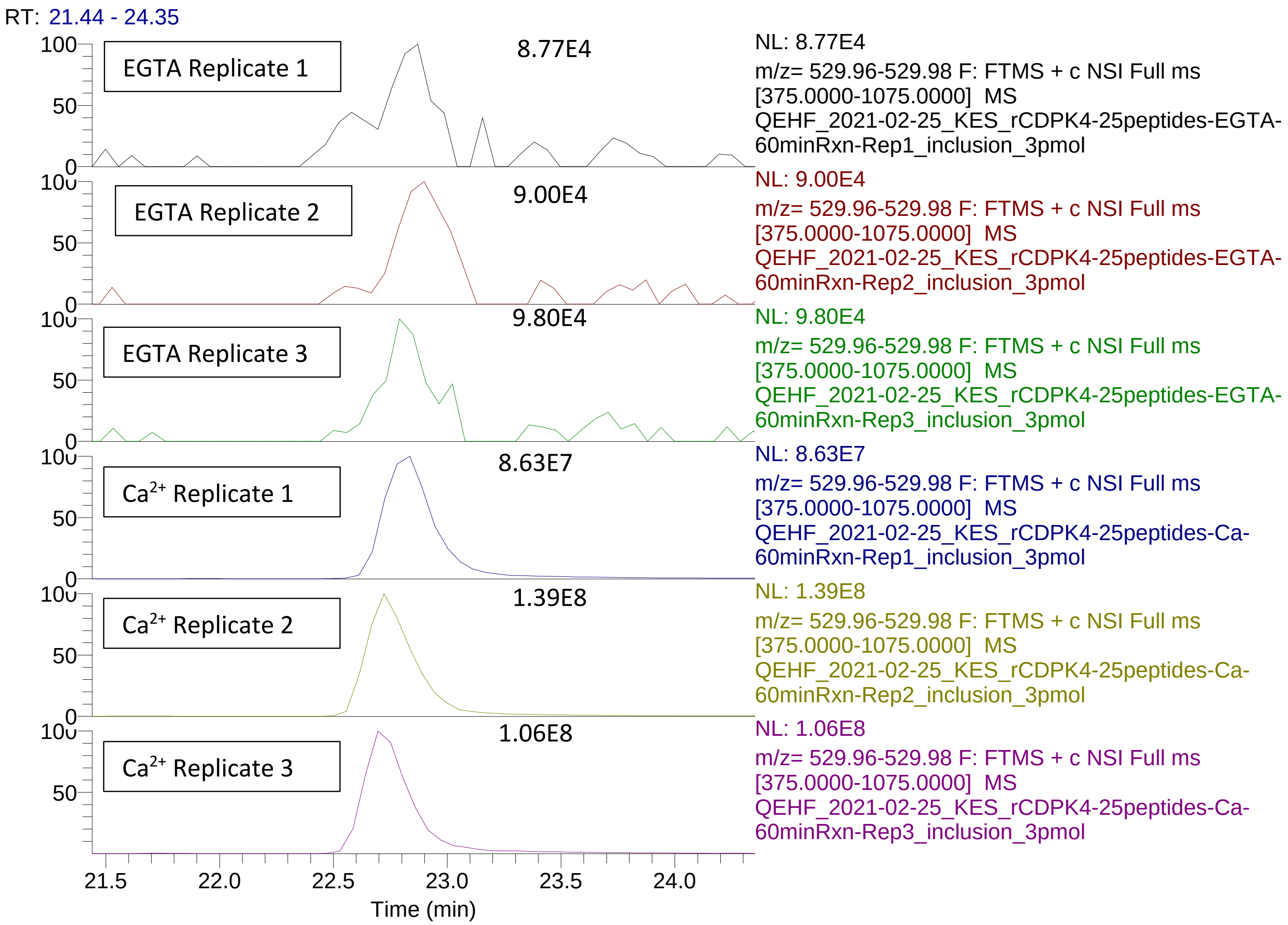


Extracted ion chromatograms for 529.97 m/z (charge +3). Peaks are labeled with peak intensity. This peptide was confidently identified in all runs as the peptide syntide 2 phosphopeptpide, PLARTLS[+79.97]VAGLPGKK. PfCDPK4 is inactive in the presence of the calcium chelator EGTA and active in the presence of $Ca^{2+}$. Phosphopeptide abundance in the presence of EGTA was near background levels, indicating near complete inactivation of the kinase. By contrast, the presence of $Ca^{2+}$ was in the kinase buffer led to near complete conversion of the peptide to phosphopeptide.

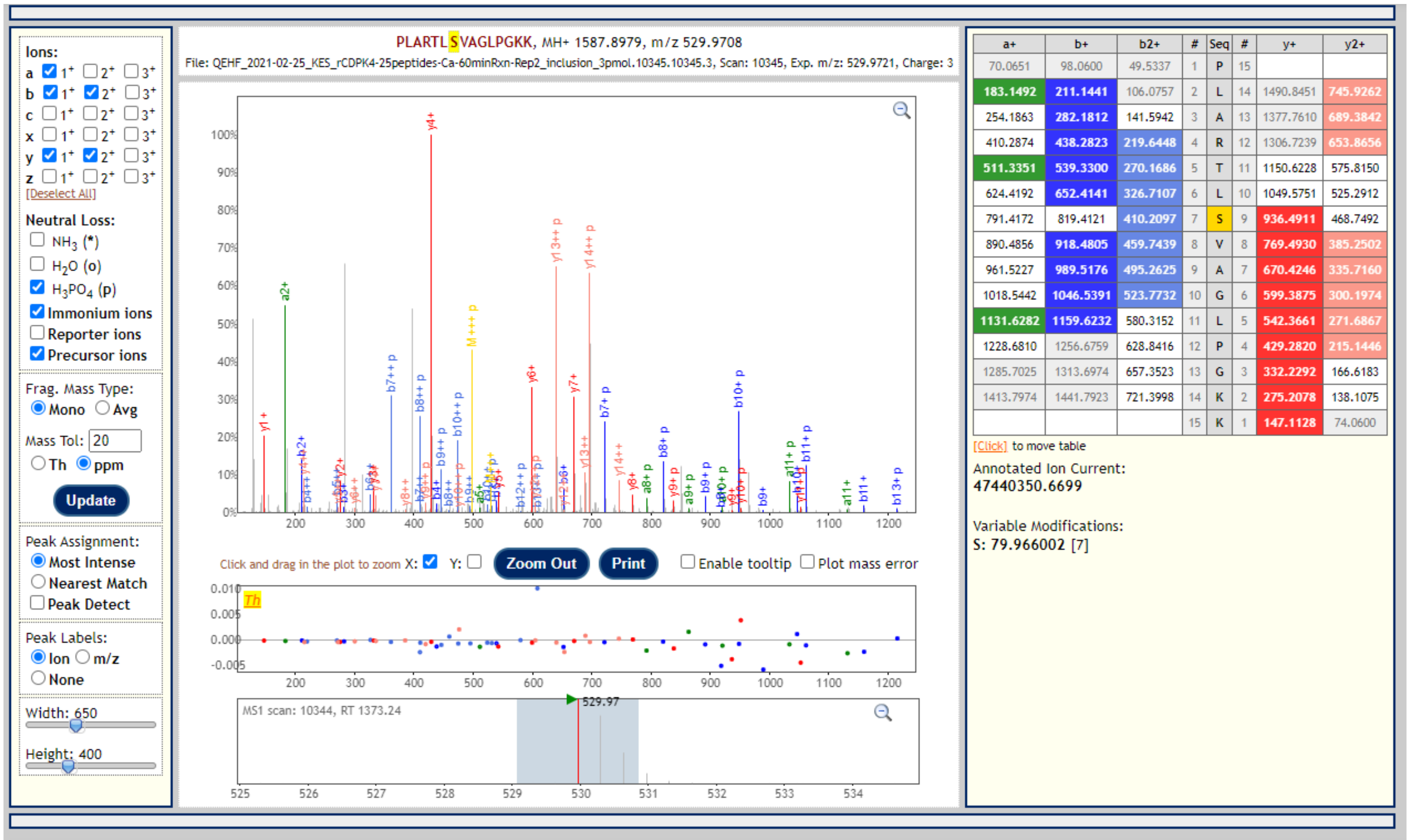


| a+ | b+ | b2+ | # | Seq | # | y+ | y2+ |
|---|---|---|---|---|---|---|---|
| 70.0651 | 98.0600 | 49.5337 | 1 | P | 15 | | |
| 183.1492 | 211.1441 | 106.0757 | 2 | L | 14 | 1490.8451 | 745.9262 |
| 254.1863 | 282.1812 | 141.5942 | 3 | A | 13 | 1377.7610 | 689.3842 |
| 410.2874 | 438.2823 | 219.6448 | 4 | R | 12 | 1306.7239 | 653.8656 |
| 511.3351 | 539.3300 | 270.1686 | 5 | T | 11 | 1150.6228 | 575.8150 |
| 624.4192 | 652.4141 | 326.7107 | 6 | L | 10 | 1049.5751 | 525.2912 |
| 791.4172 | 819.4121 | 410.2097 | 7 | S | 9 | 936.4911 | 468.7492 |
| 890.4856 | 918.4805 | 459.7439 | 8 | V | 8 | 769.4930 | 385.2502 |
| 961.5227 | 989.5176 | 495.2625 | 9 | A | 7 | 670.4246 | 335.7160 |
| 1018.5442 | 1046.5391 | 523.7732 | 10 | G | 6 | 599.3875 | 300.1974 |
| 1131.6282 | 1159.6232 | 580.3152 | 11 | L | 5 | 542.3661 | 271.6867 |
| 1228.6810 | 1256.6759 | 628.8416 | 12 | P | 4 | 429.2820 | 215.1446 |
| 1285.7025 | 1313.6974 | 657.3523 | 13 | G | 3 | 332.2292 | 166.6183 |
| 1413.7974 | 1441.7923 | 721.3998 | 14 | K | 2 | 275.2078 | 138.1075 |
| | | | 15 | K | 1 | 147.1128 | 74.0600 |

MS2 spectrum from the apex of the 529.97 m/z (3+) peak identifying the syntide 2 phosphopeptide $Ca^{2+}$ replicate 1.
PeptideProphet probability = 1.000
PTMProphet localization scores = PLART(0.012)LS(0.988)VAGLPGKK, confidently localizing the phosphosite.

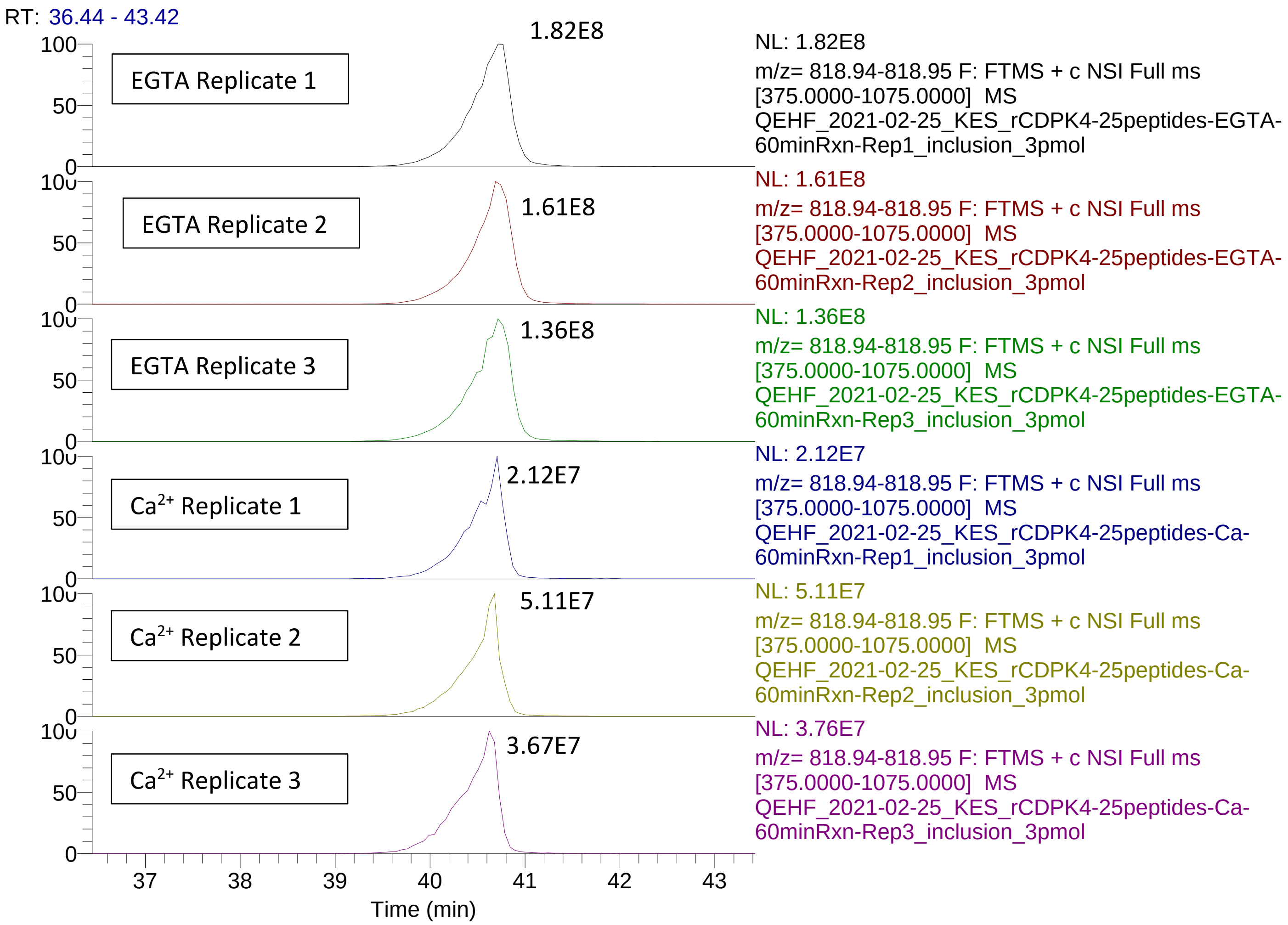


Extracted ion chromatograms for 818.95 m/z (charge +2). Peaks are labeled with peak intensity. This peptide was confidently identified in all runs as the unmodified peptide SLPYEKALSSTISQI from the uncharacterized protein PF3D7_0417600. PfCDPK4 is inactive in the presence of the calcium chelator EGTA and active in the presence of $Ca^{2+}$. In the replicates with active PfCDPK4, the peptide abundance was depleted an average of 77% (85% for the +3 ion), indicating that the peptide was phosphorylated.

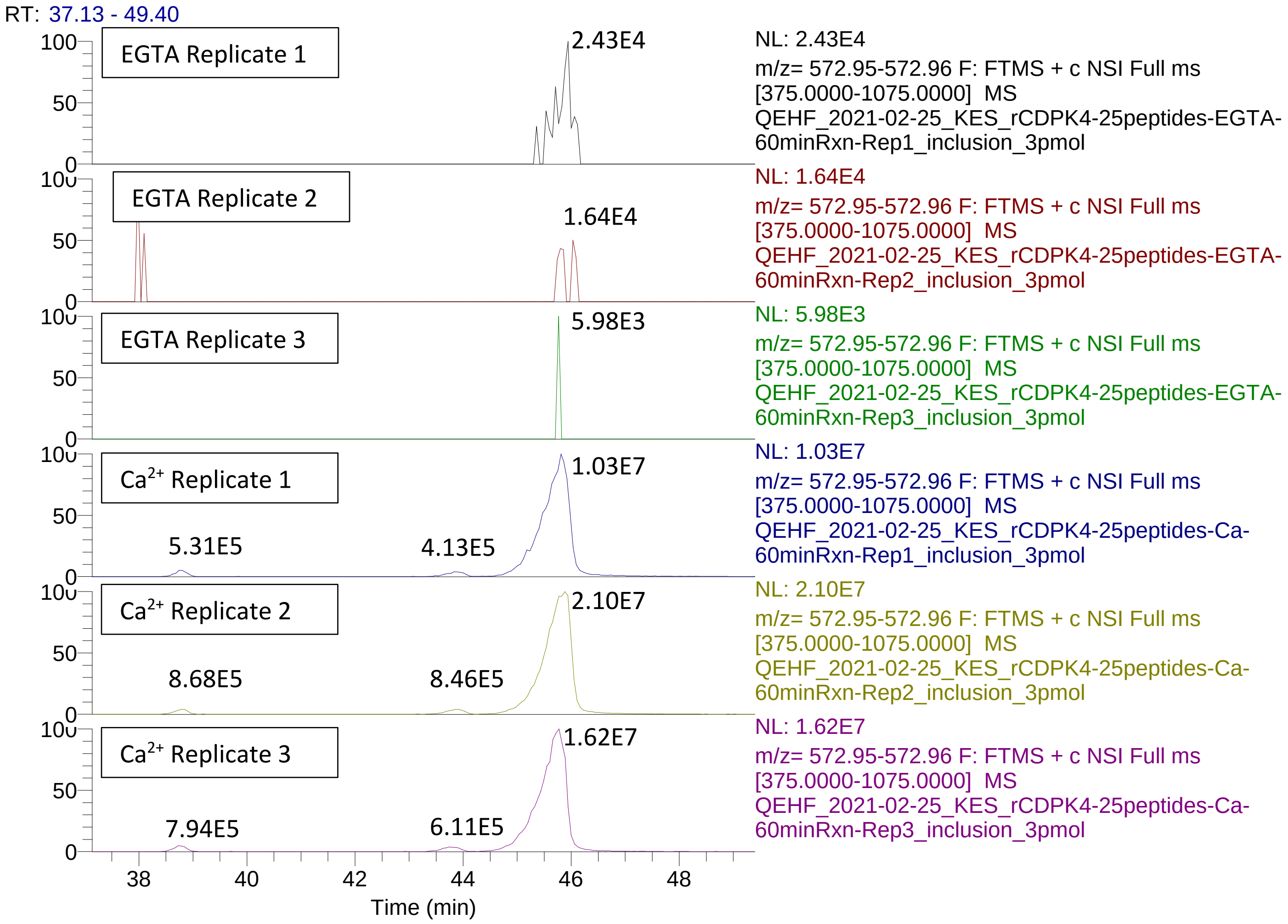


Extracted ion chromatograms for 572.95 m/z (charge +3), which matches the mass of the singly phosphorylated peptide SLPYEKALSSTISQI from the uncharacterized protein PF3D7_0417600. PfCDPK4 is inactive in the presence of the calcium chelator EGTA and active in the presence of $Ca^{2+}$. Peaks are labeled with peak intensity. In the replicates with active PfCDPK4, there are three isobaric species present at distinct retention times, with the major species at retention time 45.8 min. These likely represent three positional isomers of the phosphopeptide. The MS2 spectra from this species could not confidently determine the peptide sequence, nor could the phosphosite be confidently localized (see below). However, the fact that the unmodified peptide was depleted by over 77% in the presence of activated PfCDPK4 concomitant with appearance of a peptide matching the mass of the phosphopeptide supports the claim that the peptide was phosphorylated by PfCDPK4.

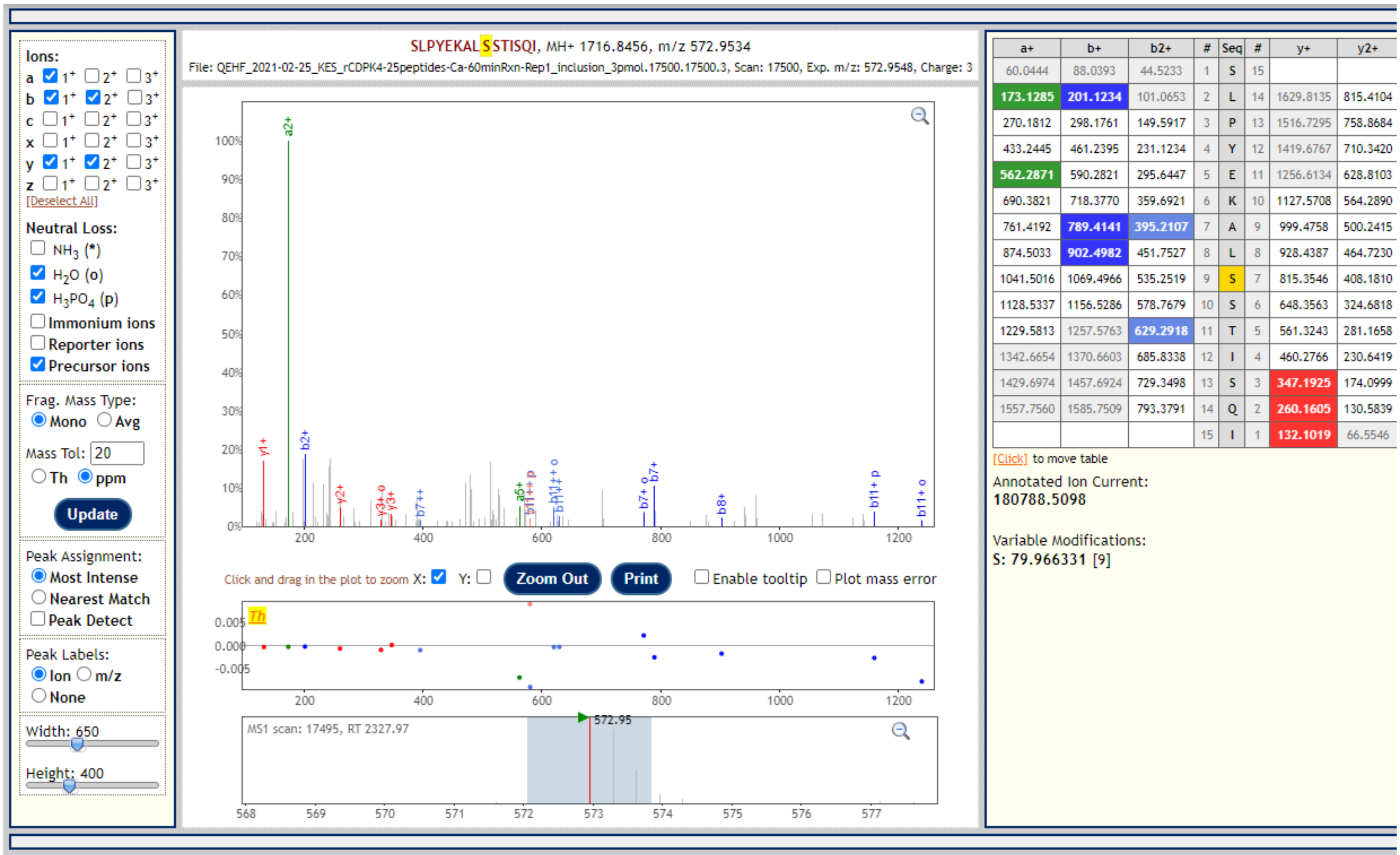


| a+ | b+ | b2+ | # | Seq | # | y+ | y2+ |
|---|---|---|---|---|---|---|---|
| 60.0444 | 88.0393 | 44.5233 | 1 | S | 15 | | |
| 173.1285 | 201.1234 | 101.0653 | 2 | L | 14 | 1629.8135 | 815.4104 |
| 270.1812 | 298.1761 | 149.5917 | 3 | P | 13 | 1516.7295 | 758.8684 |
| 433.2445 | 461.2395 | 231.1234 | 4 | Y | 12 | 1419.6767 | 710.3420 |
| 562.2871 | 590.2821 | 295.6447 | 5 | E | 11 | 1256.6134 | 628.8103 |
| 690.3821 | 718.3770 | 359.6921 | 6 | K | 10 | 1127.5708 | 564.2890 |
| 761.4192 | 789.4141 | 395.2107 | 7 | A | 9 | 999.4758 | 500.2415 |
| 874.5033 | 902.4982 | 451.7527 | 8 | L | 8 | 928.4387 | 464.7230 |
| 1041.5016 | 1069.4966 | 535.2519 | 9 | S | 7 | 815.3546 | 408.1810 |
| 1128.5337 | 1156.5286 | 578.7679 | 10 | S | 6 | 648.3563 | 324.6818 |
| 1229.5813 | 1257.5763 | 629.2918 | 11 | T | 5 | 561.3243 | 281.1658 |
| 1342.6654 | 1370.6603 | 685.8338 | 12 | I | 4 | 460.2766 | 230.6419 |
| 1429.6974 | 1457.6924 | 729.3498 | 13 | S | 3 | 347.1925 | 174.0999 |
| 1557.7560 | 1585.7509 | 793.3791 | 14 | Q | 2 | 260.1605 | 130.5839 |
| | | | 15 | I | 1 | 132.1019 | 66.5546 |

MS2 spectrum from the apex of the 572.95 m/z (3+) peak at 38.7 min in $Ca^{2+}$ replicate 1.

PeptideProphet probability = 0.7096, below the cut-off for identifying spectra.

There are insufficient peaks to localize the phosphosite.

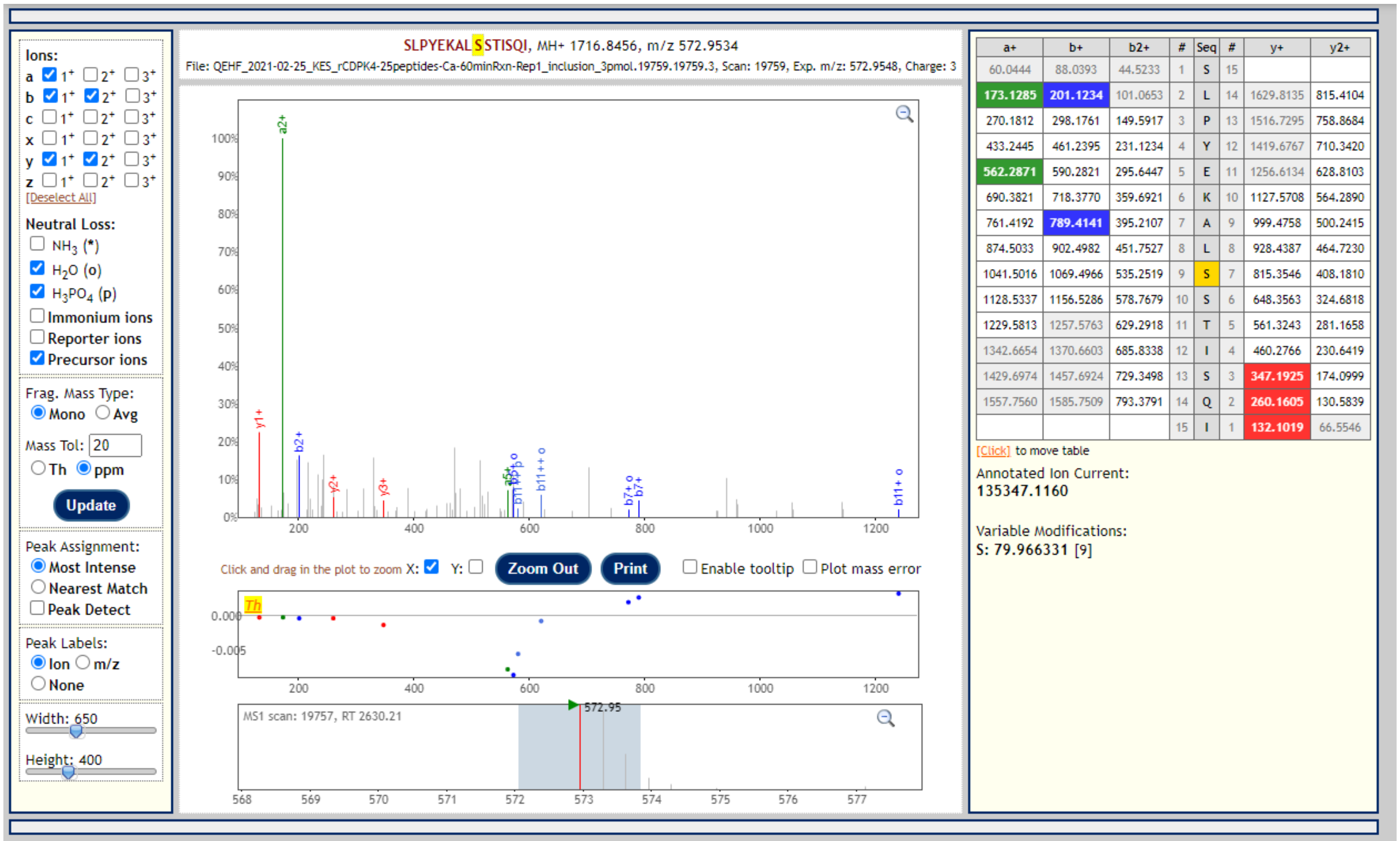


MS2 spectrum from apex of 572.95 m/z (3+) peak at 43.8 min in $Ca^{2+}$ replicate 1.
PeptideProphet probability = 0.4826, below the cutoff for identifying spectra.
There are insufficient peaks to localize the phosphosite.

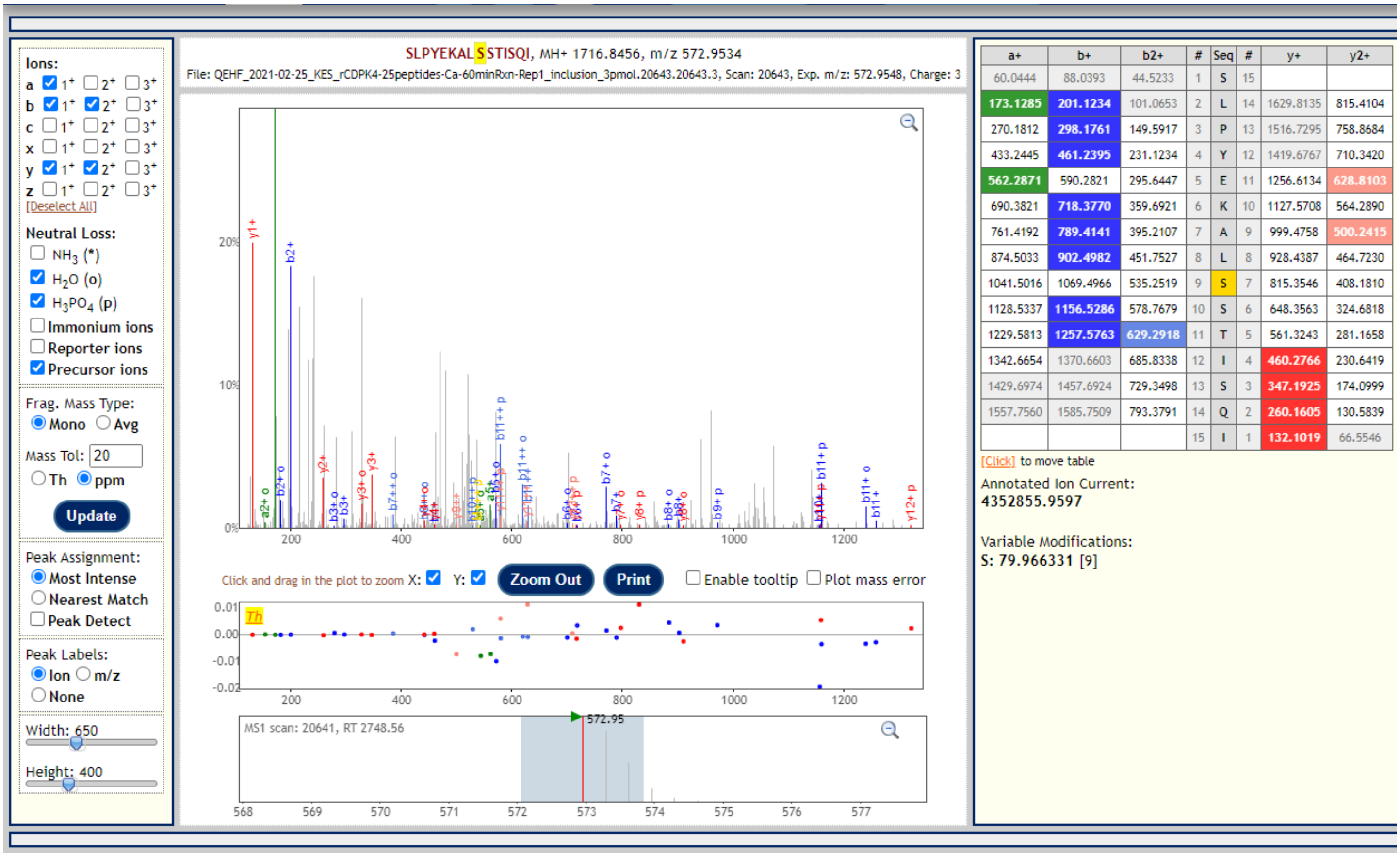


MS2 spectrum from apex of 572.95 m/z (3+) peak at 45.8 min in $Ca^{2+}$ replicate 1.
PeptideProphet probability = 0.4740, below the cutoff for identifying spectra.
There are insufficient peaks to localize the phosphosite.
