## Supplementary data S1-S6 for "*Plasmodium falciparum* Calcium Dependent Protein Kinase 4 is critical for male gametogenesis and transmission to the mosquito vector": Supplementary_Data_S5_Primers.docx

**Table S1: PCR Primers used in the studies:**

| **Oligo** | **Forward (5’-3’)** |
| --- | --- |
| PfCDPK4 5’Homo For | T**GCGGCCGC**GTTTCCCTATCTTTTCAGTGCATTTTG |
| PfCDPK4 5’Homo Rev | TTATGGTTTATTTGATAATGGTTGATACCTTCTTCTTTATATATTGTCTAATG |
| PfCDPK4 3’Homo For | ATAAAGAAGAAGGTATCAACCATTATCAAATAAACCATAAATCAATTCAATA |
| PfCDPK4 3’Homo Rev | TAA**GTCGAC**TTAAACTGTGATGATACACATTCTCACTTG |
| PfCDPK4Guide For | TATTGAAATGAAAGAGAGTAGTGT |
| PfCDPK4Guide Rev | AAACACACTACTCTCTTTCATTTC |
| PfCDPK4 Geno5 For | TTTTCTAGGTAAAGTATTAATATATTGTGTGTAAA |
| PfCDPK4 Geno5 Rev | ACACTACTCTCTTTCATTTCATGTCTCTCA |
| PfCDPK4 Geno3 For | GAAGTGGATCAAAATAATGATGGAGAA |
| PfCDPK4 Geno3 Rev | GTATTTAAATATGTGCAGCACAATTTATTC |
