## Supplementary data S1-S6 for "*Plasmodium falciparum* Calcium Dependent Protein Kinase 4 is critical for male gametogenesis and transmission to the mosquito vector": Supplementary_Data_S6_LC-MS_data_processing_parameters.pdf

#### Comet parameters used for gametocyte phosphopeptide LC-MS/MS data

```
# comet_version 2020.01 rev. 3
# Comet MS/MS search engine parameters file.
# Everything following the '#' symbol is treated as a comment.

database_name = Pfalciparum_human_cRAP_DECOY.fasta
decoy_search = 0                       # 0=no (default), 1=concatenated search, 2=separate search
peff_format = 0                        # 0=no (normal fasta, default), 1=PEFF PSI-MOD, 2=PEFF Unimod
peff_obo =                             # path to PSI Mod or Unimod OBO file

num_threads = 8                        # 0=poll CPU to set num threads; else specify num threads directly (max
128)

#
# masses
#
peptide_mass_tolerance = 10.00
peptide_mass_units = 2                 # 0=amu, 1=mmu, 2=ppm
mass_type_parent = 1                   # 0=average masses, 1=monoisotopic masses
mass_type_fragment = 1                 # 0=average masses, 1=monoisotopic masses
precursor_tolerance_type = 0           # 0=MH+ (default), 1=precursor m/z; only valid for amu/mmu tolerances
isotope_error = 3                      # 0=off, 1=0/1 (C13 error), 2=0/1/2, 3=0/1/2/3, 4=-8/-4/0/4/8 (for +4/+8
labeling)

#
# search enzyme
#
search_enzyme_number = 1               # choose from list at end of this params file
search_enzyme2_number = 0              # second enzyme; set to 0 if no second enzyme
num_enzyme_termini = 2                 # 1 (semi-digested), 2 (fully digested, default), 8 C-term unspecific , 9
N-term unspecific
allowed_missed_cleavage = 2            # maximum value is 5; for enzyme search

#
# Up to 9 variable modifications are supported
# format:  <mass> <residues> <0=variable/else binary> <max_mods_per_peptide> <term_distance> <n/c-term>
<required> <neutral_loss>
#     e.g. 79.966331 STY 0 3 -1 0 0 97.976896
#
variable_mod01 = 15.9949 M 0 3 -1 0 0 0.0
variable_mod02 = 79.966331 STY 0 3 -1 0 0 0
variable_mod03 = 42.010565 n 0 1 1 0 0 0.0          #acetylation of protein N-terminus at N-terminal residue or
next residue
variable_mod04 = 0.0 X 0 3 -1 0 0 0.0
variable_mod05 = 0.0 X 0 3 -1 0 0 0.0
variable_mod06 = 0.0 X 0 3 -1 0 0 0.0
variable_mod07 = 0.0 X 0 3 -1 0 0 0.0
variable_mod08 = 0.0 X 0 3 -1 0 0 0.0
variable_mod09 = 0.0 X 0 3 -1 0 0 0.0
max_variable_mods_in_peptide = 5
require_variable_mod = 0

#
# fragment ions
#
# ion trap ms/ms:  1.0005 tolerance, 0.4 offset (mono masses), theoretical_fragment_ions = 1
# high res ms/ms:    0.02 tolerance, 0.0 offset (mono masses), theoretical_fragment_ions = 0,
spectrum_batch_size = 15000
#
fragment_bin_tol = 0.02              # binning to use on fragment ions
fragment_bin_offset = 0.0              # offset position to start the binning (0.0 to 1.0)
theoretical_fragment_ions = 0          # 0=use flanking peaks, 1=M peak only
use_A_ions = 0
use_B_ions = 1
use_C_ions = 0
use_X_ions = 0
use_Y_ions = 1
use_Z_ions = 0
use_Z1_ions = 0
use_NL_ions = 0                        # 0=no, 1=yes to consider NH3/H2O neutral loss peaks
```

```
#
# output
#
output_sqtfile = 0                     # 0=no, 1=yes  write sqt file
output_txtfile = 0                     # 0=no, 1=yes  write tab-delimited txt file
output_pepxmlfile = 1                  # 0=no, 1=yes  write pepXML file
output_mzidentmlfile = 0               # 0=no, 1=yes  write mzIdentML file
output_percolatorfile = 0              # 0=no, 1=yes  write Percolator pin file
print_expect_score = 1                 # 0=no, 1=yes to replace Sp with expect in out & sqt
num_output_lines = 5                   # num peptide results to show

sample_enzyme_number = 1               # Sample enzyme which is possibly different than the one applied to the
search.
                                       # Used to calculate NTT & NMC in pepXML output (default=1 for trypsin).

#
# mzXML parameters
#
scan_range = 0 0                       # start and end scan range to search; either entry can be set
independently
precursor_charge = 0 0                 # precursor charge range to analyze; does not override any existing
charge; 0 as 1st entry ignores parameter
override_charge = 0                    # 0=no, 1=override precursor charge states, 2=ignore precursor charges
outside precursor_charge range, 3=see online
ms_level = 2                           # MS level to analyze, valid are levels 2 (default) or 3
activation_method = ALL                # activation method; used if activation method set; allowed ALL, CID,
ECD, ETD, ETD+SA, PQD, HCD, IRMPD, SID

#
# misc parameters
#
digest_mass_range = 600.0 5000.0       # MH+ peptide mass range to analyze
peptide_length_range = 5 63            # minimum and maximum peptide length to analyze (default 1 63; max length
63)
num_results = 100                      # number of search hits to store internally
max_duplicate_proteins = 20            # maximum number of additional duplicate protein names to report for each
peptide ID; -1 reports all duplicates
max_fragment_charge = 3                # set maximum fragment charge state to analyze (allowed max 5)
max_precursor_charge = 5               # set maximum precursor charge state to analyze (allowed max 9)
nucleotide_reading_frame = 0           # 0=proteinDB, 1-6, 7=forward three, 8=reverse three, 9=all six
clip_nterm_methionine = 0              # 0=leave sequences as-is; 1=also consider sequence w/o N-term methionine
spectrum_batch_size = 15000            # max. # of spectra to search at a time; 0 to search the entire scan
range in one loop
decoy_prefix = DECOY_                  # decoy entries are denoted by this string which is pre-pended to each
protein accession
equal_I_and_L = 1                      # 0=treat I and L as different; 1=treat I and L as same
output_suffix =                        # add a suffix to output base names i.e. suffix "-C" generates base-
C.pep.xml from base.mzXML input
mass_offsets =                         # one or more mass offsets to search (values substracted from
deconvoluted precursor mass)
precursor_NL_ions =                    # one or more precursor neutral loss masses, will be added to xcorr
analysis

#
# spectral processing
#
minimum_peaks = 10                     # required minimum number of peaks in spectrum to search (default 10)
minimum_intensity = 0                  # minimum intensity value to read in
remove_precursor_peak = 0              # 0=no, 1=yes, 2=all charge reduced precursor peaks (for ETD),
3=phosphate neutral loss peaks
remove_precursor_tolerance = 1.5       # +- Da tolerance for precursor removal
clear_mz_range = 0.0 0.0               # for iTRAQ/TMT type data; will clear out all peaks in the specified m/z
range

#
# additional modifications
#

add_Cterm_peptide = 0.0
add_Nterm_peptide = 0.0
add_Cterm_protein = 0.0
add_Nterm_protein = 0.0
```

```
add_G_glycine = 0.0000                 # added to G - avg.  57.0513, mono.  57.02146
add_A_alanine = 0.0000                 # added to A - avg.  71.0779, mono.  71.03711
add_S_serine = 0.0000                  # added to S - avg.  87.0773, mono.  87.03203
add_P_proline = 0.0000                 # added to P - avg.  97.1152, mono.  97.05276
add_V_valine = 0.0000                  # added to V - avg.  99.1311, mono.  99.06841
add_T_threonine = 0.0000               # added to T - avg. 101.1038, mono. 101.04768
add_C_cysteine = 57.021464             # added to C - avg. 103.1429, mono. 103.00918
add_L_leucine = 0.0000                 # added to L - avg. 113.1576, mono. 113.08406
add_I_isoleucine = 0.0000              # added to I - avg. 113.1576, mono. 113.08406
add_N_asparagine = 0.0000              # added to N - avg. 114.1026, mono. 114.04293
add_D_aspartic_acid = 0.0000           # added to D - avg. 115.0874, mono. 115.02694
add_Q_glutamine = 0.0000               # added to Q - avg. 128.1292, mono. 128.05858
add_K_lysine = 0.0000                  # added to K - avg. 128.1723, mono. 128.09496
add_E_glutamic_acid = 0.0000           # added to E - avg. 129.1140, mono. 129.04259
add_M_methionine = 0.0000              # added to M - avg. 131.1961, mono. 131.04048
add_H_histidine = 0.0000               # added to H - avg. 137.1393, mono. 137.05891
add_F_phenylalanine = 0.0000           # added to F - avg. 147.1739, mono. 147.06841
add_U_selenocysteine = 0.0000          # added to U - avg. 150.0379, mono. 150.95363
add_R_arginine = 0.0000                # added to R - avg. 156.1857, mono. 156.10111
add_Y_tyrosine = 0.0000                # added to Y - avg. 163.0633, mono. 163.06333
add_W_tryptophan = 0.0000              # added to W - avg. 186.0793, mono. 186.07931
add_O_pyrrolysine = 0.0000             # added to O - avg. 237.2982, mono  237.14773
add_B_user_amino_acid = 0.0000         # added to B - avg.   0.0000, mono.   0.00000
add_J_user_amino_acid = 0.0000         # added to J - avg.   0.0000, mono.   0.00000
add_X_user_amino_acid = 0.0000         # added to X - avg.   0.0000, mono.   0.00000
add_Z_user_amino_acid = 0.0000         # added to Z - avg.   0.0000, mono.   0.00000

#
# COMET_ENZYME_INFO _must_ be at the end of this parameters file
#
[COMET_ENZYME_INFO]
0.  Cut_everywhere         0      -           -
1.  Trypsin                1      KR          P
2.  Trypsin/P              1      KR          -
3.  Lys_C                  1      K           P
4.  Lys_N                  0      K           -
5.  Arg_C                  1      R           P
6.  Asp_N                  0      D           -
7.  CNBr                   1      M           -
8.  Glu_C                  1      DE          P
9.  PepsinA                1      FL          P
10. Chymotrypsin           1      FWYL        P
```

### Trans-Proteomic Pipeline parameters used for gametocyte phosphopeptide LC-MS/MS data

```
>InteractParser <WT|KO>.interact.pep.xml LUMOS_2020-09-29_KES_PfGams_<WT|KO>_*.pep.xml -I

>RefreshParser <WT|KO>.interact.pep.xml `DatabaseParser <WT|KO>.interact.pep.xml`

>PeptideProphetParser <WT|KO>.interact.pep.xml ZACCMASS PPM MINPROB=0 DECOY=DECOY NONPARAM

>RefreshParser <WT|KO>.interact.pep.xml `DatabaseParser <WT|KO>.interact.pep.xml`

>InterProphetParser THREADS=8 <WT|KO>.interact.pep.xml <WT|KO>.interact.iproph.pep.xml

>PTMProphetParser MAXTHREADS=8 MINPROB=0.9 STY:79.966:-97.97690,M:15.9949 STATIC FRAGPPMTOL=15
<WT|KO>.interact.iproph.pep.xml <WT|KO>.interact.ptm.iproph.pep.xml
```

#### Comet parameters used for *in vitro* kinase assay of synthetic peptides LC-MS/MS data

```
# comet_version 2020.01 rev. 3
# Comet MS/MS search engine parameters file.
# Everything following the '#' symbol is treated as a comment.

database_name = Peptides.fasta
decoy_search = 1                       # 0=no (default), 1=concatenated search, 2=separate search
peff_format = 0                        # 0=no (normal fasta, default), 1=PEFF PSI-MOD, 2=PEFF Unimod
peff_obo =                             # path to PSI Mod or Unimod OBO file

num_threads = 8                        # 0=poll CPU to set num threads; else specify num threads directly (max
128)

#
# masses
#
peptide_mass_tolerance = 10
peptide_mass_units = 2                 # 0=amu, 1=mmu, 2=ppm
mass_type_parent = 1                   # 0=average masses, 1=monoisotopic masses
mass_type_fragment = 1                 # 0=average masses, 1=monoisotopic masses
precursor_tolerance_type = 0           # 0=MH+ (default), 1=precursor m/z; only valid for amu/mmu tolerances
isotope_error = 0                      # 0=off, 1=0/1 (C13 error), 2=0/1/2, 3=0/1/2/3, 4=-8/-4/0/4/8 (for +4/+8
labeling)

#
# search enzyme
#
search_enzyme_number = 1               # choose from list at end of this params file
search_enzyme2_number = 0              # second enzyme; set to 0 if no second enzyme
num_enzyme_termini = 2                 # 1 (semi-digested), 2 (fully digested, default), 8 C-term unspecific , 9
N-term unspecific
allowed_missed_cleavage = 5            # maximum value is 5; for enzyme search

#
# Up to 9 variable modifications are supported
# format:  <mass> <residues> <0=variable/else binary> <max_mods_per_peptide> <term_distance> <n/c-term>
<required> <neutral_loss>
#     e.g. 79.966331 STY 0 3 -1 0 0 97.976896
#
variable_mod01 = 15.9949 M 0 3 -1 0 0 0.0
variable_mod02 = 79.966331 STY 0 3 -1 0 0 0
variable_mod03 = 0.0 X 0 3 -1 0 0 0.0
variable_mod04 = 0.0 X 0 3 -1 0 0 0.0
variable_mod05 = 0.0 X 0 3 -1 0 0 0.0
variable_mod06 = 0.0 X 0 3 -1 0 0 0.0
variable_mod07 = 0.0 X 0 3 -1 0 0 0.0
variable_mod08 = 0.0 X 0 3 -1 0 0 0.0
variable_mod09 = 0.0 X 0 3 -1 0 0 0.0
max_variable_mods_in_peptide = 5
require_variable_mod = 0

#
# fragment ions
#
# ion trap ms/ms:  1.0005 tolerance, 0.4 offset (mono masses), theoretical_fragment_ions = 1
# high res ms/ms:    0.02 tolerance, 0.0 offset (mono masses), theoretical_fragment_ions = 0,
spectrum_batch_size = 15000
#
fragment_bin_tol = 0.02              # binning to use on fragment ions
fragment_bin_offset = 0.0              # offset position to start the binning (0.0 to 1.0)
theoretical_fragment_ions = 0          # 0=use flanking peaks, 1=M peak only
use_A_ions = 0
use_B_ions = 1
use_C_ions = 0
use_X_ions = 0
use_Y_ions = 1
use_Z_ions = 0
use_Z1_ions = 0
use_NL_ions = 0                        # 0=no, 1=yes to consider NH3/H2O neutral loss peaks

#
# output
#
```

```
output_sqtfile = 0                     # 0=no, 1=yes  write sqt file
output_txtfile = 0                     # 0=no, 1=yes  write tab-delimited txt file
output_pepxmlfile = 1                  # 0=no, 1=yes  write pepXML file
output_mzidentmlfile = 0               # 0=no, 1=yes  write mzIdentML file
output_percolatorfile = 0              # 0=no, 1=yes  write Percolator pin file
print_expect_score = 1                 # 0=no, 1=yes to replace Sp with expect in out & sqt
num_output_lines = 5                   # num peptide results to show

sample_enzyme_number = 1               # Sample enzyme which is possibly different than the one applied to the
search.
                                       # Used to calculate NTT & NMC in pepXML output (default=1 for trypsin).

#
# mzXML parameters
#
scan_range = 0 0                       # start and end scan range to search; either entry can be set
independently
precursor_charge = 0 0                 # precursor charge range to analyze; does not override any existing
charge; 0 as 1st entry ignores parameter
override_charge = 0                    # 0=no, 1=override precursor charge states, 2=ignore precursor charges
outside precursor_charge range, 3=see online
ms_level = 2                           # MS level to analyze, valid are levels 2 (default) or 3
activation_method = ALL                # activation method; used if activation method set; allowed ALL, CID,
ECD, ETD, ETD+SA, PQD, HCD, IRMPD, SID

#
# misc parameters
#
digest_mass_range = 600.0 5000.0       # MH+ peptide mass range to analyze
peptide_length_range = 5 63            # minimum and maximum peptide length to analyze (default 1 63; max length
63)
num_results = 100                      # number of search hits to store internally
max_duplicate_proteins = 20            # maximum number of additional duplicate protein names to report for each
peptide ID; -1 reports all duplicates
max_fragment_charge = 3                # set maximum fragment charge state to analyze (allowed max 5)
max_precursor_charge = 4               # set maximum precursor charge state to analyze (allowed max 9)
nucleotide_reading_frame = 0           # 0=proteinDB, 1-6, 7=forward three, 8=reverse three, 9=all six
clip_nterm_methionine = 0              # 0=leave sequences as-is; 1=also consider sequence w/o N-term methionine
spectrum_batch_size = 15000            # max. # of spectra to search at a time; 0 to search the entire scan
range in one loop
decoy_prefix = DECOY_                  # decoy entries are denoted by this string which is pre-pended to each
protein accession
equal_I_and_L = 1                      # 0=treat I and L as different; 1=treat I and L as same
output_suffix =                        # add a suffix to output base names i.e. suffix "-C" generates base-
C.pep.xml from base.mzXML input
mass_offsets =                         # one or more mass offsets to search (values substracted from
deconvoluted precursor mass)
precursor_NL_ions =                    # one or more precursor neutral loss masses, will be added to xcorr
analysis

#
# spectral processing
#
minimum_peaks = 10                     # required minimum number of peaks in spectrum to search (default 10)
minimum_intensity = 0                  # minimum intensity value to read in
remove_precursor_peak = 0              # 0=no, 1=yes, 2=all charge reduced precursor peaks (for ETD),
3=phosphate neutral loss peaks
remove_precursor_tolerance = 1.5       # +- Da tolerance for precursor removal
clear_mz_range = 0.0 0.0               # for iTRAQ/TMT type data; will clear out all peaks in the specified m/z
range

#
# additional modifications
#

add_Cterm_peptide = 0.0
add_Nterm_peptide = 0.0
add_Cterm_protein = 0.0
add_Nterm_protein = 0.0

add_G_glycine = 0.0000                 # added to G - avg.  57.0513, mono.  57.02146
add_A_alanine = 0.0000                 # added to A - avg.  71.0779, mono.  71.03711
add_S_serine = 0.0000                  # added to S - avg.  87.0773, mono.  87.03203
```

```
add_P_proline = 0.0000                 # added to P - avg.  97.1152, mono.  97.05276
add_V_valine = 0.0000                  # added to V - avg.  99.1311, mono.  99.06841
add_T_threonine = 0.0000               # added to T - avg. 101.1038, mono. 101.04768
add_C_cysteine = 0.0000                # added to C - avg. 103.1429, mono. 103.00918
add_L_leucine = 0.0000                 # added to L - avg. 113.1576, mono. 113.08406
add_I_isoleucine = 0.0000              # added to I - avg. 113.1576, mono. 113.08406
add_N_asparagine = 0.0000              # added to N - avg. 114.1026, mono. 114.04293
add_D_aspartic_acid = 0.0000           # added to D - avg. 115.0874, mono. 115.02694
add_Q_glutamine = 0.0000               # added to Q - avg. 128.1292, mono. 128.05858
add_K_lysine = 0.0000                  # added to K - avg. 128.1723, mono. 128.09496
add_E_glutamic_acid = 0.0000           # added to E - avg. 129.1140, mono. 129.04259
add_M_methionine = 0.0000              # added to M - avg. 131.1961, mono. 131.04048
add_H_histidine = 0.0000               # added to H - avg. 137.1393, mono. 137.05891
add_F_phenylalanine = 0.0000           # added to F - avg. 147.1739, mono. 147.06841
add_U_selenocysteine = 0.0000          # added to U - avg. 150.0379, mono. 150.95363
add_R_arginine = 0.0000                # added to R - avg. 156.1857, mono. 156.10111
add_Y_tyrosine = 0.0000                # added to Y - avg. 163.0633, mono. 163.06333
add_W_tryptophan = 0.0000              # added to W - avg. 186.0793, mono. 186.07931
add_O_pyrrolysine = 0.0000             # added to O - avg. 237.2982, mono  237.14773
add_B_user_amino_acid = 0.0000         # added to B - avg.   0.0000, mono.   0.00000
add_J_user_amino_acid = 0.0000         # added to J - avg.   0.0000, mono.   0.00000
add_X_user_amino_acid = 0.0000         # added to X - avg.   0.0000, mono.   0.00000
add_Z_user_amino_acid = 0.0000         # added to Z - avg.   0.0000, mono.   0.00000

#
# COMET_ENZYME_INFO _must_ be at the end of this parameters file
#
[COMET_ENZYME_INFO]
0.  Cut_everywhere         0      -           -
1.  Trypsin                1      KR          ACDEFGHIKLMNPQRSTVWY
2.  Trypsin/P              1      KR          -
3.  Lys_C                  1      K           P
4.  Lys_N                  0      K           -
5.  Arg_C                  1      R           P
6.  Asp_N                  0      D           -
7.  CNBr                   1      M           -
8.  Glu_C                  1      DE          P
9.  PepsinA                1      FL          P
10. Chymotrypsin           1      FWYL        P
```

#### Trans-Proteomic Pipeline parameters used for *in vitro* kinase assay of synthetic peptides  LC-MS/MS data

```
>InteractParser interact.pep.xml *.pep.xml -I

>RefreshParser interact.pep.xml `DatabaseParser interact.pep.xml`

>PeptideProphetParser interact.pep.xml ZACCMASS PPM MINPROB=0 DECOYPROBS EXPECTSCORE DECOY=DECOY NONPARAM NONMC
NONTT

>RefreshParser interact.pep.xml `DatabaseParser interact.pep.xml`

>cp interact.pep.xml interact.XPRESS.pep.xml

>XPressPeptideParser interact.XPRESS.pep.xml -l -m10 -a -c5 -p1
#-l label free -m10 mass tolerance -a use ppm -c5 minimum 5 points per peak (default) -p1 include 1 C13 isotope
peak (default)
```

### Comet parameters used for recombinant PfCDPK4 autophosphorylation LC-MS/MS data

```
# comet_version 2020.01 rev. 3
# Comet MS/MS search engine parameters file.
# Everything following the '#' symbol is treated as a comment.

database_name = rPfCDPK4-MBP_Ecoli_cRAP_DECOY.fasta
decoy_search = 0                        # 0=no (default), 1=concatenated search, 2=separate search
peff_format = 0                         # 0=no (normal fasta, default), 1=PEFF PSI-MOD, 2=PEFF Unimod
peff_obo =                              # path to PSI Mod or Unimod OBO file

num_threads = 0                         # 0=poll CPU to set num threads; else specify num threads directly (max
128)

#
# masses
#
peptide_mass_tolerance = 10.00
peptide_mass_units = 2                  # 0=amu, 1=mmu, 2=ppm
mass_type_parent = 1                    # 0=average masses, 1=monoisotopic masses
mass_type_fragment = 1                  # 0=average masses, 1=monoisotopic masses
precursor_tolerance_type = 0            # 0=MH+ (default), 1=precursor m/z; only valid for amu/mmu tolerances
isotope_error = 3                       # 0=off, 1=0/1 (C13 error), 2=0/1/2, 3=0/1/2/3, 4=-8/-4/0/4/8 (for +4/+8
labeling)

#
# search enzyme
#
search_enzyme_number = 1                # choose from list at end of this params file
search_enzyme2_number = 0               # second enzyme; set to 0 if no second enzyme
num_enzyme_termini = 1                  # 1 (semi-digested), 2 (fully digested, default), 8 C-term unspecific , 9
N-term unspecific
allowed_missed_cleavage = 2             # maximum value is 5; for enzyme search

#
# Up to 9 variable modifications are supported
# format:  <mass> <residues> <0=variable/else binary> <max_mods_per_peptide> <term_distance> <n/c-term>
<required> <neutral_loss>
#     e.g. 79.966331 STY 0 3 -1 0 0 97.976896
#
variable_mod01 = 79.966331 STY 0 3 -1 0 0 0
variable_mod02 = 0.0 X 0 3 -1 0 0 0.0
variable_mod03 = 0.0 X 0 3 -1 0 0 0.0
variable_mod04 = 0.0 X 0 3 -1 0 0 0.0
variable_mod05 = 0.0 X 0 3 -1 0 0 0.0
variable_mod06 = 0.0 X 0 3 -1 0 0 0.0
variable_mod07 = 0.0 X 0 3 -1 0 0 0.0
variable_mod08 = 0.0 X 0 3 -1 0 0 0.0
variable_mod09 = 0.0 X 0 3 -1 0 0 0.0
max_variable_mods_in_peptide = 5
require_variable_mod = 0

#
# fragment ions
#
# ion trap ms/ms:  1.0005 tolerance, 0.4 offset (mono masses), theoretical_fragment_ions = 1
# high res ms/ms:    0.02 tolerance, 0.0 offset (mono masses), theoretical_fragment_ions = 0,
spectrum_batch_size = 15000
#
fragment_bin_tol = 0.02               # binning to use on fragment ions
fragment_bin_offset = 0.0               # offset position to start the binning (0.0 to 1.0)
theoretical_fragment_ions = 0           # 0=use flanking peaks, 1=M peak only
use_A_ions = 0
use_B_ions = 1
use_C_ions = 0
use_X_ions = 0
use_Y_ions = 1
use_Z_ions = 0
use_Z1_ions = 0
use_NL_ions = 0                         # 0=no, 1=yes to consider NH3/H2O neutral loss peaks

#
# output
#
```

```
output_sqtfile = 0                     # 0=no, 1=yes  write sqt file
output_txtfile = 0                     # 0=no, 1=yes  write tab-delimited txt file
output_pepxmlfile = 1                  # 0=no, 1=yes  write pepXML file
output_mzidentmlfile = 0               # 0=no, 1=yes  write mzIdentML file
output_percolatorfile = 0              # 0=no, 1=yes  write Percolator pin file
print_expect_score = 1                 # 0=no, 1=yes to replace Sp with expect in out & sqt
num_output_lines = 5                   # num peptide results to show

sample_enzyme_number = 1               # Sample enzyme which is possibly different than the one applied to the
search.
                                       # Used to calculate NTT & NMC in pepXML output (default=1 for trypsin).

#
# mzXML parameters
#
scan_range = 0 0                       # start and end scan range to search; either entry can be set
independently
precursor_charge = 0 0                 # precursor charge range to analyze; does not override any existing
charge; 0 as 1st entry ignores parameter
override_charge = 0                    # 0=no, 1=override precursor charge states, 2=ignore precursor charges
outside precursor_charge range, 3=see online
ms_level = 2                           # MS level to analyze, valid are levels 2 (default) or 3
activation_method = ALL                # activation method; used if activation method set; allowed ALL, CID,
ECD, ETD, ETD+SA, PQD, HCD, IRMPD, SID

#
# misc parameters
#
digest_mass_range = 600.0 5000.0       # MH+ peptide mass range to analyze
peptide_length_range = 5 63            # minimum and maximum peptide length to analyze (default 1 63; max length
63)
num_results = 100                      # number of search hits to store internally
max_duplicate_proteins = 20            # maximum number of additional duplicate protein names to report for each
peptide ID; -1 reports all duplicates
max_fragment_charge = 3                # set maximum fragment charge state to analyze (allowed max 5)
max_precursor_charge = 6               # set maximum precursor charge state to analyze (allowed max 9)
nucleotide_reading_frame = 0           # 0=proteinDB, 1-6, 7=forward three, 8=reverse three, 9=all six
clip_nterm_methionine = 0              # 0=leave sequences as-is; 1=also consider sequence w/o N-term methionine
spectrum_batch_size = 15000            # max. # of spectra to search at a time; 0 to search the entire scan
range in one loop
decoy_prefix = DECOY_                  # decoy entries are denoted by this string which is pre-pended to each
protein accession
equal_I_and_L = 1                      # 0=treat I and L as different; 1=treat I and L as same
output_suffix =                        # add a suffix to output base names i.e. suffix "-C" generates base-
C.pep.xml from base.mzXML input
mass_offsets =                         # one or more mass offsets to search (values substracted from
deconvoluted precursor mass)
precursor_NL_ions =                    # one or more precursor neutral loss masses, will be added to xcorr
analysis

#
# spectral processing
#
minimum_peaks = 10                     # required minimum number of peaks in spectrum to search (default 10)
minimum_intensity = 0                  # minimum intensity value to read in
remove_precursor_peak = 0              # 0=no, 1=yes, 2=all charge reduced precursor peaks (for ETD),
3=phosphate neutral loss peaks
remove_precursor_tolerance = 1.5       # +- Da tolerance for precursor removal
clear_mz_range = 0.0 0.0               # for iTRAQ/TMT type data; will clear out all peaks in the specified m/z
range

#
# additional modifications
#

add_Cterm_peptide = 0.0
add_Nterm_peptide = 0.0
add_Cterm_protein = 0.0
add_Nterm_protein = 0.0

add_G_glycine = 0.0000                 # added to G - avg.  57.0513, mono.  57.02146
add_A_alanine = 0.0000                 # added to A - avg.  71.0779, mono.  71.03711
add_S_serine = 0.0000                  # added to S - avg.  87.0773, mono.  87.03203
```

```
add_P_proline = 0.0000                 # added to P - avg.  97.1152, mono.  97.05276
add_V_valine = 0.0000                  # added to V - avg.  99.1311, mono.  99.06841
add_T_threonine = 0.0000               # added to T - avg. 101.1038, mono. 101.04768
add_C_cysteine = 57.021464             # added to C - avg. 103.1429, mono. 103.00918
add_L_leucine = 0.0000                 # added to L - avg. 113.1576, mono. 113.08406
add_I_isoleucine = 0.0000              # added to I - avg. 113.1576, mono. 113.08406
add_N_asparagine = 0.0000              # added to N - avg. 114.1026, mono. 114.04293
add_D_aspartic_acid = 0.0000           # added to D - avg. 115.0874, mono. 115.02694
add_Q_glutamine = 0.0000               # added to Q - avg. 128.1292, mono. 128.05858
add_K_lysine = 0.0000                  # added to K - avg. 128.1723, mono. 128.09496
add_E_glutamic_acid = 0.0000           # added to E - avg. 129.1140, mono. 129.04259
add_M_methionine = 0.0000              # added to M - avg. 131.1961, mono. 131.04048
add_H_histidine = 0.0000               # added to H - avg. 137.1393, mono. 137.05891
add_F_phenylalanine = 0.0000           # added to F - avg. 147.1739, mono. 147.06841
add_U_selenocysteine = 0.0000          # added to U - avg. 150.0379, mono. 150.95363
add_R_arginine = 0.0000                # added to R - avg. 156.1857, mono. 156.10111
add_Y_tyrosine = 0.0000                # added to Y - avg. 163.0633, mono. 163.06333
add_W_tryptophan = 0.0000              # added to W - avg. 186.0793, mono. 186.07931
add_O_pyrrolysine = 0.0000             # added to O - avg. 237.2982, mono  237.14773
add_B_user_amino_acid = 0.0000         # added to B - avg.   0.0000, mono.   0.00000
add_J_user_amino_acid = 0.0000         # added to J - avg.   0.0000, mono.   0.00000
add_X_user_amino_acid = 0.0000         # added to X - avg.   0.0000, mono.   0.00000
add_Z_user_amino_acid = 0.0000         # added to Z - avg.   0.0000, mono.   0.00000

#
# COMET_ENZYME_INFO _must_ be at the end of this parameters file
#
[COMET_ENZYME_INFO]
0.  Cut_everywhere         0      -           -
1.  Trypsin                1      KR          P
2.  Trypsin/P              1      KR          -
3.  Lys_C                  1      K           P
4.  Lys_N                  0      K           -
5.  Arg_C                  1      R           P
6.  Asp_N                  0      D           -
7.  CNBr                   1      M           -
8.  Glu_C                  1      DE          P
9.  PepsinA                1      FL          P
10. Chymotrypsin           1      FWYL        P
```

#### Trans-Proteomic Pipeline parameters used for recombinant PfCDPK4 autophosphorylation LC-MS/MS data

```
>InteractParser interact.pep.xml *.pep.xml -I

>RefreshParser interact.pep.xml `DatabaseParser interact.pep.xml`

>PeptideProphetParser interact.pep.xml ZACCMASS PPM MINPROB=0 DECOYPROBS DECOY=DECOY EXPECTSCORE NONPARAM

>RefreshParser interact.pep.xml `DatabaseParser interact.pep.xml`

>PTMProphetParser MAXTHREADS=8 MINPROB=0.5 STY:79.966:-97.97690 STATIC FRAGPPMTOL=15 interact.pep.xml
interact.ptm.pep.xml
```
